# Demonstration of local adaptation of maize landraces by reciprocal transplantation

**DOI:** 10.1101/2021.03.25.437076

**Authors:** Garrett M. Janzen, María Rocío Aguilar-Rangel, Carolina Cíntora-Martínez, Karla Azucena Blöcher-Juárez, Eric González-Segovia, Anthony J. Studer, Daniel E. Runcie, Sherry A. Flint-Garcia, Rubén Rellán-Álvarez, Ruairidh J. H. Sawers, Matthew B. Hufford

**Affiliations:** Department of Ecology, Evolution, and Organismal Biology, Iowa State University, Ames, Iowa, USA 50011; Department of Plant Biology, University of Georgia, Athens, Georgia, USA 30602; Langebio, Cinvestav, Km 9.6 Libramiento Norte Carretera Len, Irapuato, Guanajuato, Mexico 36821; Department of Crop Sciences, University of Illinois Urbana-Champaign, 1201 West Gregory Drive, Urbana, Illinois, USA 61801; Department of Plant Sciences, University of California-Davis, 278 Robbins, Berkeley, California, USA 95616; Agricultural Research Service, United States Department of Agriculture, Columbia, Missouri, 65211; University of Missouri, 301 Curtis Hall, Columbia, Missouri, USA 65211; Molecular and Structural Biochemistry, North Carolina State University, 128 Polk Hall, Raleigh, North Carolina, USA 27695-7622; Department of Plant Science, Pennsylvania State University, University Park, Pennsylvania, USA, 16802

**Keywords:** *Zea mays*, landrace, population genetics, local adaptation, highland adaptation, reciprocal transplant

## Abstract

Populations are locally adapted when they exhibit higher fitness than foreign populations in their native habitat. Maize landrace adaptations to highland and lowland conditions are of interest to researchers and breeders. To determine the prevalence and strength of local adaptation in maize landraces, we performed a reciprocal transplant experiment across an elevational gradient in Mexico. We grew 120 landraces, grouped into four populations (Mexican Highland, Mexican Lowland, South American Highland, South American Lowland), in Mexican highland and lowland common gardens and collected phenotypes relevant to fitness, as well as reported highland-adaptive traits such as anthocyanin pigmentation and macrohair density. 67k DArTseq markers were generated from field specimens to allow comparison between phenotypic patterns and population genetic structure.

We found phenotypic patterns consistent with local adaptation, though these patterns differ between the Mexican and South American populations. While population genetic structure largely recapitulates drift during post-domestication dispersal, landrace phenotypes reflect adaptations to native elevation. Quantitative trait *Q_ST_* was greater than neutral *F_ST_* for many traits, signaling divergent directional selection between pairs of populations. All populations exhibited higher fitness metric values when grown at their native elevation, and Mexican landraces had higher fitness than South American landraces when grown in our Mexican sites. Highland populations expressed generally higher anthocyanin pigmentation than lowland populations, and more so in the highland site than in the lowland site. Macrohair density was largely non-plastic, and Mexican landraces and highland landraces were generally more pilose. Analysis of *δ*^13^C indicated that lowland populations may have lower WUE. Each population demonstrated garden-specific correlations between highland trait expression and fitness, with stronger positive correlations in the highland site.

These results give substance to the long-held presumption of local adaptation of New World maize landraces to elevation and other environmental variables across North and South America.

## 1 Introduction

Populations evolve adaptations to selective pressures imparted by biotic and abiotic environments. Over time, given sufficiently low genetic drift and gene flow, theory predicts that a population will adapt to the particular selective pressures of its local environment (Leimu and Fischer, 2008). In particular, populations are said to be locally adapted when they meet the “Foreign vs. Local” criterion of local adaptation, in which a local population exhibits higher fitness than foreign populations when grown in the same environment (Kawecki and Ebert, 2004).

Traditionally, attempts to identify and quantify local adaptation in natural populations have relied on common garden experiments (Turesson, 1922; Clausen et al., 1940; Fraser et al., 2011; Savolainen et al., 2013). Reciprocal transplant experiments are in many cases preferable to common garden experiments, as the scale, complexity, and variety of the environments of the included populations can be modeled more holistically, rather than being reduced to single or few environmental variables (Kawecki and Ebert, 2004; Savolainen et al., 2013; Limpens et al., 2012; Gibson et al., 2016). Exposing individuals from different populations to common environments can reveal that environments affect populations differently, a situation known as genotype-by-environment (*G* × *E*) interaction (Savolainen et al., 2013). Local adaptation is a type of *G* × *E* interaction in which a population has higher fitness in its native environment than any other non-native population in that environment. Local adaptation is illustrated by crossing fitness reaction norms in a reciprocal transplant experiment (Kawecki and Ebert, 2004; Savolainen et al., 2013).

Maize (Zea *mays* subsp. *mays*) is an extensively studied model system of high agronomic (Shiferaw et al., 2011), economic (Shiferaw et al., 2011; Ranum et al., 2014), cultural (Fernandez Suarez et al., 2013; Perales, 2016), and scientific (Dumas and Mogensen, 1993; Fedoroff, 2001; Stern et al., 2004) value. Maize was domesticated in the lowlands of the Balsas River Valley in Mexico from the teosinte taxon *Zea mays* subsp. *parviglumis* roughly 9000 years BP (Matsuoka et al., 2002). From there, maize was carried across North America and into South America as early as 6000 years BP (Grobman et al., 2012; Bush et al., 1989), north into the present-day United States by about 4500 years BP (Merrill et al., 2009), and around the world as part of the Columbian exchange (Tenaillon and Charcosset, 2011; Van Heerwaarden et al., 2011). Presently, maize is grown across a greater range of elevations and latitudes than any other crop (Ruìz Corral et al., 2008; Shiferaw et al., 2011), experiencing a broad range of temperature, precipitation, and soil types.

At locations along the historical range expansion of maize, farmers selected lines that were both suitable for growth in their local environment and desirable for human consumption and applications. Over generations of propagation and selection, this process formed varietal populations called landraces. These landraces are grown and maintained by smallholder farmers to the present day as dynamic, evolving populations (Mijangos-Cortes et al., 2007; Dyer and López-Feldman, 2013) with low but significant gene flow between them (Ortega, 1995) (see Villa et al. (2005) for a review of the defining characteristics of landraces). Most of the arable land in Mexico is managed by subsistence farms that cultivate maize landraces (Bellon et al., 2018). Landraces are typically out-yielded by modern hybrids in industrial agricultural contexts, but in their own home environments, landraces can and often do out-perform commercial hybrids (Bellon et al., 2018; Perales, 2016; Bellon et al., 2003; Mercer and Perales, 2018).

Maize landraces exhibit diverse morphological, physiological, and phenological characteristics, many of which covary with climate, soil type and quality, and geography (Wellhausen EJ, 1952). While farmers consciously select primarily for ear characteristics that are indirectly related to survival and reproduction (kernel filling, ear size, varietal consistency (Louette and Smale, 2000; Prasanna et al., 2010)), the environment selects for plant survival and reproduction (Cleveland and Soleri, 2007). The combination of these selective factors comprise the agroecosystem to which landraces adapt (Villa et al., 2005; Bracco et al., 2012).

Some of the most striking adaptations in maize landraces are in response to elevation (Eagles and Lothrop, 1994). Highland conditions present challenges for maize survival and productivity. At higher elevation, the atmosphere is thinner, leading to colder temperatures and less filtering of solar radiation. Marked phenotypic variation and genetic structure are correlated with elevation, though elevation itself may not be the causal agent (Dyer and López-Feldman, 2013). In at least some high-elevation regions in Mexico, adaptations are hypothesized to be imparted via introgression from the maize wild relative *Zea mays* subsp. *mexicana* (hereafter *“mexicana*”), which is adapted to cool, dry highland conditions (Lauter et al., 2004; Hufford et al., 2012; Janzen et al., 2018; Rodríguez-Zapata et al., 2021). Notable similarities between highland maize and *mexicana* include highly pigmented and pilose leaf sheaths (Doebley, 1984). Hufford et al. (2013) found that *mexicana* introgression into sympatric maize in Mexico overlapped chromosomal regions identified as QTL by Lauter et al. (2004) for pilosity and pigmentation (though other loci influence variance in these traits, e.g. *b1*, Selinger and Chandler (2001)). Dark red pigments absorb solar radiation, warming the plant. Pilosity increases surface friction, which decreases wind speed across the surface of the plant. This boundary layer around the plant reduces both heat loss and transpiration which can be advantageous in cool, dry regions (Schuepp, 1993; Chalker-Scott, 1999).

There are multiple reasons to suspect that the nature of highland adaptations may differ significantly between landrace populations and between highland regions. First, highland adaptation seems to have evolved mostly independently in Mesoamerica and South America. Takuno et al. (2015) found that highland landraces in Mexico and South America were independently derived from lowland germplasm through selection on standing variation and *de novo* mutations, with little genomic evidence of convergent evolution. This hypothesis is supported by the absence of *mexicana* haplotypes (which are common in highland Mexican landraces and lacking in lowland Mexican landraces) in Andean highland landraces (Wang et al., 2017). Though more recent research (Wang et al., 2020) since Takuno et al. (2015) has found low but significant parallel highland adaptation between Mesoamerican and South American highland populations, which may be conferred through bi-directional human-mediated migration (Kistler et al., 2020), the predominant pattern of highland adaptation remains independent. Second, selective pressures imparted by highland (and lowland) environments in Mesoamerica and South America are not identical. The strength and direction of correlations between elevation and climatic conditions can vary from one highland region to another. Precipitation and temperature correlate with elevation differently between Mexico and South America, and between lowland habitats west and east of highland ranges. In general, across Mexico, lowland conditions range from tropical to temperate, whereas highland conditions are cooler and drier (Medina et al., 1998). In South America, eastern lowlands neighbor the Amazon Basin, western coastal regions are arid, and southern highlands and lowlands become drier with increasing distance from the equatorial tropics (Sarmiento, 1975). The Andean rain shadow produces geographic regions with elevational gradients of cooler, moister highlands and hotter, dryer lowlands, across which indigenous farmers continue to cultivate maize and other crops (Brush, 1976). Because precipitation and temperature do not uniformly correlate with elevation, landraces that have evolved adaptations to high-elevation bioclimatic conditions in South America may be ill-suited for conditions found at the same elevation in Mexico.

A better assessment of maize landrace local adaptation may prove valuable for modern maize breeders. The intense breeding programs that have developed modern inbred lines have drawn from limited germplasm and, through selection, have further reduced genetic diversity and capacity for adaptive plasticity (Gage et al., 2017). Reincorporation of landrace germplasm can restore key genetic variants that impart adaptations to challenging environments. Despite this potential, and despite a number of studies that report that local adaptation is pervasive among maize landraces (Harlan, 1975; Villa et al., 2005; Navarro et al., 2017; Bracco et al., 2012), research has not fully addressed whether maize landraces broadly do, in fact, exhibit reciprocal home-site advantage, the definition of local adaptation. Landrace geographical extents have been shown to correspond to elevational and climatic factors (Ruìz Corral et al., 2008; Arteaga et al., 2016; Aguirre-Liguori et al., 2019), supporting (but not demonstrating) local adaptation. Reciprocal transplant experiments set along an elevational gradient in the Mexican state of Chiapas (Mercer et al., 2008; Mercer and Perales, 2018) have shown that landraces local to that area exhibit local adaptation. Taking a different approach, a recent study by Gates et al. (2019) found that landrace F1 hybrids (landrace individuals crossed with locally-adapted testers) exhibit higher fitness and yield when grown at common garden sites closer to the native elevation of the landrace parent. This research identified promising candidate local adaptation loci among landraces and provides strong evidence of local adaptation. However, as this study utilized hybrids from landraces with only limited sampling outside Mexico, it does not necessarily demonstrate landrace local adaptation in a larger context. The extent of local adaptation among maize landraces, therefore, has not been fully established.

To investigate the extent and degree of local adaptation between highland and lowland maize landraces, we conducted an elevational reciprocal transplant experiment. We compared fitness metrics and reportedly highland-adaptive traits (macrohair and anthocyanin pigmentation) from highland and lowland Mexican and South American maize landrace populations grown in highland and lowland Mexican sites to investigate differential plastic responses to highland conditions. We also compared quantitative trait differentiation (*Q_ST_*) to neutral genetic variance between populations (*F_ST_*) to find traits under divergent directional selection, and correlated values of highland-adaptive traits with fitness traits to investigate their elevation-specific relationship with fitness.

## 2 Methods

### 2.1 Field Experiment Design

Landrace accessions from CIMMYT that met the following criteria were considered for inclusion in this experiment:

1. Accessions are present in the Seeds of Discovery (SeeDs) dataset (Pixley et al., 2017).
2. Accessions had latitude and longitude data from North or South America.
3. The elevation of the accession was from below 1000 m or above 2000 m.

From eligible accessions, 30 pairs of highland and lowland accessions were chosen from both Mexico and South America (120 accessions total) such that both landraces of a pair were collected from the same 1-degree of latitude bin, and all pairwise distances between accessions were greater than 50 km. These 120 samples were split into four populations (Mexican Highland, Mexican Lowland, South American Highland, South American Lowland, hereafter “Mex High,” “Mex Low,” “SA High,” and “SA Low”) with 30 accessions per population. We note that our provisional population designations are designed to reflect continental and elevational distinctions and not necessarily population genetic structure, and that we use the word “Mexican” to refer the North American populations despite the fact that two of the accessions are from Guatemala.

The two common garden sites that comprise this reciprocal transplant are the Winter Services nursery site near Puerto Vallarta in the Pacific coastal lowlands (elevation 54 m) of Mexico (hereafter “Low Site”), and a CIMMYT field site near the town of Metepec in the highlands (elevation 2852 m) of the Mexican Central Plateau (hereafter “High Site”). Seed lines were regenerated at the field site for one generation prior to the experiment to reduce seed storage and maternal effects. Best local practices for irrigation, fertilizer, and pest/weed control were used at both sites. The High Site field experiment was conducted in the summer of 2016. The Low Site field experiment was conducted in the winter of 2016, but virus damage led us to repeat the field experiment at the same site in the winter of 2017. Certain traits were collected from both years of the Low Site. A map of the field sites and geographical origin of each accession and boxplots summarizing the elevational and annual precipitation distributions of these four populations are presented in Figure 1.

**Figure 1:**
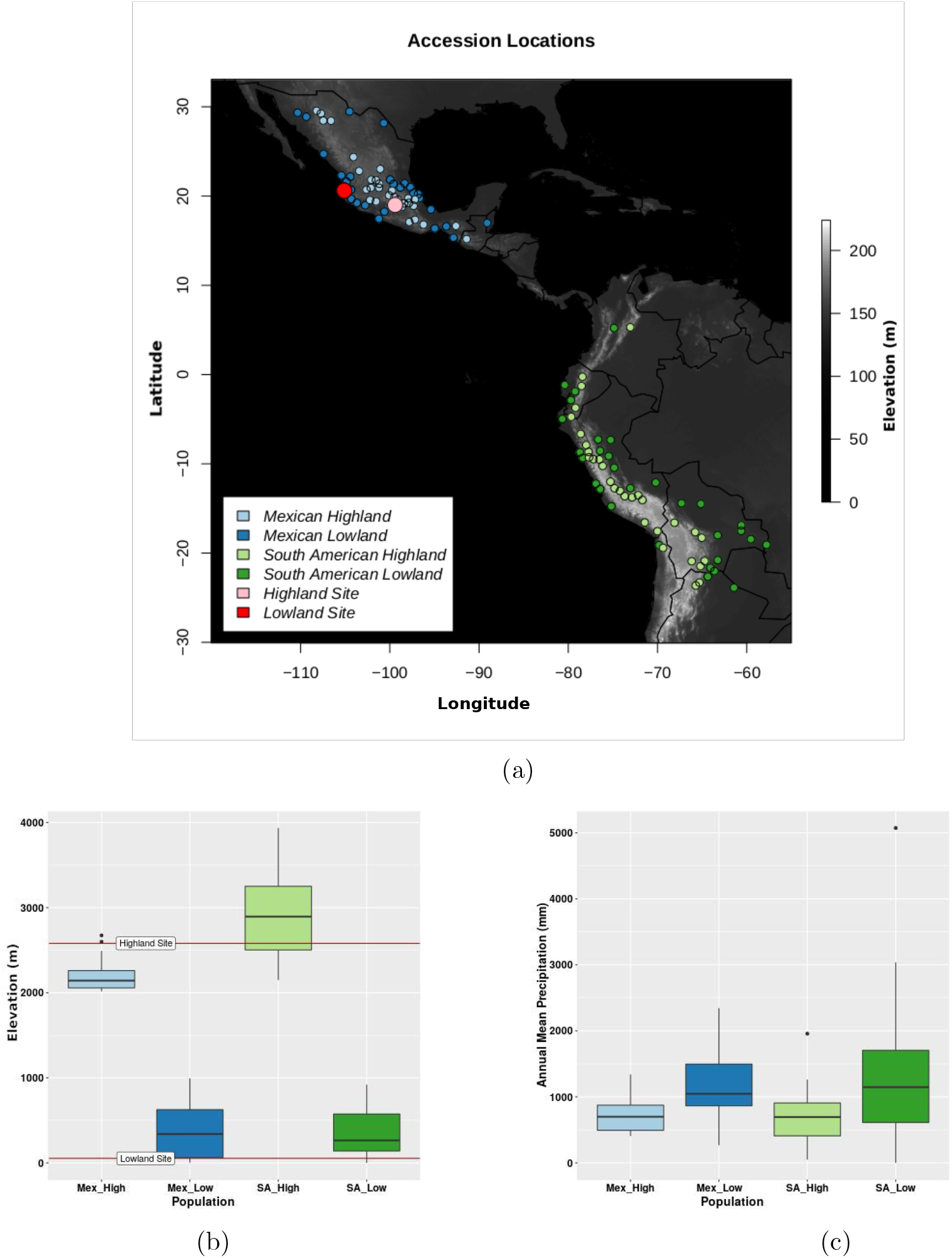
Geography and climate of 120 landraces and common garden sites. (1a) Location of collection sites of landraces and common garden sites. (1b) Elevation of collection sites of landraces. Red lines indicate the elevations of the highland and lowland common garden sites. (1c) Annual mean precipitation of collection sites of landraces.

Each field was arranged in a complete block design with two blocks of 120 rows of 15 seeds of a landrace accession. Landraces from latitudinal pairs were planted in adjacent rows.

### 2.2 Phenotypic and Genotypic Data Collection

Phenotypes (Table 1) were collected from the High Site and both years of the Low Site common gardens. Ear traits from the Low Site were collected from the 2016 season, but all other traits were taken from the 2017 growing season. Two healthy, representative plants from the interior of each row were selected and tagged. Individual plant phenotype data (plant height, ear height, ear number, tassel length, and tassel branch number) were collected from tagged plants. Other traits (stand count, ear-producing stand count, barrenness, and flowering time) were collected at the row level. Days to anthesis and days to silking were recorded as the number of days until 50% of the row exhibited silk emergence or anther exertion on more than half of the main tassel spike, respectively. Anthesis-silking interval is calculated as the difference in these two values.

**Table 1:** Names and descriptions of all collected phenotypes

| Code | Trait Name | Unit of Measurement | Trait Description | Level |
| --- | --- | --- | --- | --- |
| STD | Stand Count | count | Number of plants surviving to sexual maturity | Row |
| PE | Ear-Producing Stand Count | count | Number of plants surviving to produce ears | Row |
| BRN | Barrenness | 1-(PE/STD) | Percent of plants in family that produce no ears | Row |
| DTA | Days to Anthesis | count | Number of days between planting and 50% of plants exhibiting anthesis | Row |
| DTS | Days to Silking | count | Number of days between planting and 50% of plants in the row silking | Row |
| ASI | Anthesis/Silking Interval | DTS-DTA | Number of days between 50% silking and 50% anthesis | Row |
| PH | Plant Height | cm | Distance between the ground and the ligule of the flag leaf | Plant |
| EH | Ear Height | cm | Distance between the ground and the primary (top) ear-bearing node | Plant |
| TL | Tassel Length | cm | Distance from top tip of the main spike to the attachment point of the bottom branch | Plant |
| TBN | Tassel Branch Number | count | Number of tassel branches that attach to main spike | Plant |
| EN | Ear Number | count | Number of seed-producing ears produced | Plant |
| EW | Ear Weight | g | Mass of the primary ear | Plant |
| EL | Ear Length | cm | Length of the primary ear | Plant |
| KPR | Kernels per Row | count | Number of kernels in a row on the primary ear | Plant |
| ED | Ear Diameter | cm | Diameter of the primary ear | Plant |
| $\delta^{13}\text{C}$ | $\delta^{13}\text{C}$ | $[(R_{\text{Sample}}/R_{\text{Standard}}) - 1] * 1000$ | Degree of inclusion of $^{13}\text{C}$ in flag leaf tissue | Plant |
| FITplant | Agronomic Plant Fitness | $PE/15 * \sqrt{EN} * EW$ | Adjusted plant fitness | Plant |
| FITplantveg | Vegetative Plant Fitness | $PE/15 * \sqrt{EN}$ | Adjusted plant fitness | Plant |
| P_INTsolid | Pigment Intensity (Solid Pattern) | visual 0-4 code scale | Visual assessment of the intensity of anthocyanin pigmentation | Plant |
| P_INTspot | Pigment Intensity (Spot Pattern) | visual 0-4 code scale | Visual assessment of the intensity of anthocyanin pigmentation | Plant |
| P_EXTsolid | Pigment Extent (Solid Pattern) | visual % code scale | Visual assessment of the extent of anthocyanin pigmentation from the ground up | Plant |
| P_EXTspot | Pigment Extent (Spot Pattern) | visual % code scale | Visual assessment of the extent of anthocyanin pigmentation from the ground up | Plant |
| M_DENsolid | Macrohairs Density (Sheath) | visual 0-4 code scale | Visual assessment of the density of sheath macrohairs of the second leaf from top | Plant |
| M_DENmarg | Macrohairs Density (Sheath Margin) | visual 0-4 code scale | Visual assessment of the density of sheath margin macrohairs of the second leaf from top | Plant |

Primary ears from tagged plants from the High Site and the 2016 Low Site were returned to the lab to be photographed and processed for analysis. Total ear weight, ear length, ear diameter, and number of kernels per ear row were measured.

Methods for field visual assessment of anthocyanin pigmentation and macrohair were derived with modification from Lauter et al. (2004). Pigment was scored for pattern, intensity, and extent. The extent of leaf sheath anthocyanin pigmentation was visually scored as a percentage of the plant from ground level up (at 25% intervals). The intensity of leaf sheath pigmentation across the plant was visually scored on a scale of 0-4. Though all pigmentation patterns share some degree of genetic and environmental control, spots and banded/streaked patterns frequently co-occur as an induced response to pathogenic stress (Selinger and Chandler, 1999), whereas uniform pigmentation (and leaf sheath macrohair expression) is shown to be inducible by highland conditions in some landraces (particularly those harboring introgressed QTL from *mexicana*). For these reasons, the “solid” pattern may have a stronger association with highland adaptation, and other patterns may represent stress responses to other biotic and/or abiotic factors. Plants were given the categorical qualitative label of either “banded,” “spotted,” “uniform,” or “no pattern” (either no pigment present, or irregular pigment pattern). Plants with patterns of “banded” or “spotted” were binned into a “spot” group. Plants with pigment patterns “solid” and “no pattern” were binned into the group “solid.” When a plant exhibited multiple patterns, the highest-priority category was selected (uniform, then banded, then spotted, then no pattern). Macrohair density on the second leaf sheath from the top of the plant was visually scored on a scale of 0-4. Pubescence along the leaf sheath and pubescence restricted to the sheath margin may be under different genetic control, and may play different roles in highland adaptation. Therefore, plants were grouped by macrohair trait pattern (leaf sheath vs. leaf sheath margin).

Two adjusted fitness metrics were computed from the combination of several fitness traits (adapted from Mercer et al. (2008)). Agronomic plant fitness (FITplant) incorporates the count of ear-producing plants in the row (PE), the number of ears produced per plant (EN), and primary ear weight (EW). Ear-producing stand count is divided by the number of seeds planted per row (15) to produce percent survival to sexual maturity, and ear number is square-root transformed to account for diminishing yield returns of secondary, tertiary, and subsequent ears. To calculate adjusted fitness for plants that either did not produce ears by the time of harvest or were not harvested for collection of ear traits, a second plant fitness trait, vegetative plant fitness (FITplantveg), disregards ear weight from the equation. We calculate these adjusted fitness metrics thusly:

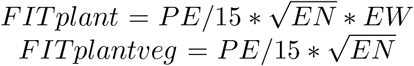

Flag leaves from tagged plants from High and 2016 Low sites were collected for Carbon isotope discrimination analysis, which was carried out at the University of Illinois (Twohey III et al., 2019). Carbon isotopic composition *δ*^13^C was calculated in reference to the international standard, Vienna Pee Dee Belemnite. The equation for *δ*^13^C (Schwarcz and Schoeninger, 1991) is as follows:

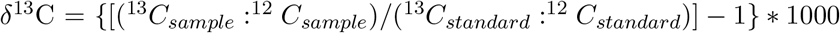

Leaf tissue samples were collected from a subset of 92 landraces in both High and Low Sites. DNA was extracted and sent to CIMMYT for DArTseq genotyping (Wenzl et al., 2004). Over 67,000 DArTseq SNP markers were generated.

### 2.3 Statistical Analyses

#### 2.3.1 *G* × *E* Interactions

We used a linear mixed-effects model (R package `lme4` (Bates et al., 2014a)) to test for phenotypic differences between landraces from each of the four populations in each trait, and how these differences changed between the two common gardens. The full model was specified as:

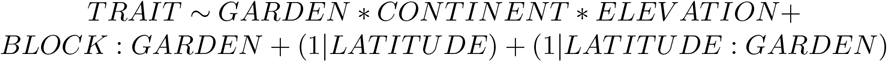

The formula calls as fixed effects GARDEN (Low Site or High Site), CONTINENT (Mexico or South America), ELEVATION (High or Low), all interaction combinations therein, BLOCK nested in GARDEN, and calls as random effects with random intercept accession LATITUDE (continuous variable) and LATITUDE/GARDEN interaction. The significance of specific treatment effects was evaluated using the `lmerTest` (Kuznetsova et al., 2017) and `lsmeans` (Lenth, 2012) R packages.

We compared each population’s phenotypes between field sites (to quantify *G* × *E* interactions), highland and lowland populations from the same continent within each field site (to quantify highland-lowland adaptation), and Mexican and South American populations from the same elevation within each field site (to quantify adaptation to continent-specific factors).

#### 2.3.2 Phenotype:Phenotype Correlations

Principal Components Analysis (function prcomp, R package `stats` (R Core Team, 2019)) was used to study the relatedness between phenotypic patterns. Data were normalized via centering and scaling. Yield traits and *δ*^13^C were available only from the second year of the Low Site, and so were excluded from PCA.

To determine the elevation-dependent fitness consequences of putatively highland-adaptive traits, we calculated Pearson correlations between fitness (FITplantveg) and traits previously identified and frequently reported as highland adaptive (Doebley, 1984; Eagles and Lothrop, 1994). FITplantveg was used rather than FITplant because FITplantveg had more complete data. These correlations were determined independently for each population in each common garden site. Additionally, the magnitude and direction of differences in fitness/highland-adaptive trait correlation coefficients between sites are taken as evidence of the trait’s adaptive role at high or low elevation. Two classes of pigment and macrohair patterns (either “solid”/”spotted” or “solid”/”margin”) were also considered separately.

#### 2.3.3 Population Genetic Relatedness

Axes of population structure were estimated from SNP data with Principal Components Analysis (R package `KRIS` (Chaichoompu et al., 2018)).

To better understand the evolutionary history between and within these four pre-defined populations, each was further sub-divided into northern and southern sub-populations (*n* = eight continent/elevation/latitude subpopulations). Pairwise Euclidean allele frequency distances between the four original populations and between the eight subpopulations were calculated (function gl.dist.pop, R package `dartR` (Gruber et al., 2018)). Population graphs (Dyer and Nason, 2004) were used to provide a graph theoretic interpretation of genetic structure between these eight subpopulations (R package `popgraph` (Dyer, 2014)).

#### 2.3.4 *Q_ST_/F_ST_* Comparison

Quantitative trait divergence (*Q_ST_*) was contrasted to the distribution of *F_ST_* for neutral genetic markers (Whitlock, 2008). For traits in which *Q_ST_* > *F_ST_*, trait divergence is greater than neutral expectations, which may be caused by directional selection (Leinonen et al., 2013).

A linear mixed effects model was used to partition phenotypic variance between population, landrace accession line, and garden/block.

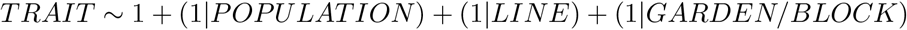

Pairwise *F_ST_* was calculated with the R function fst.each.snp.hudson (R package `dartR` (Gruber et al., 2018)). Within-population and between-population variances were calculated with the R function VarCorr (R package `lme4` (Bates et al., 2014b)), and were used to calculate *Q_ST_* following the equation below:

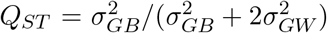

in which 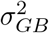 and 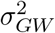 are the between- and within-population genetic variance components, respectively (Leinonen et al., 2013). Population contrasts of interest were all highland vs. all lowland, all Mexican vs. all South American, Mexican Highland vs. Mexican Lowland, and South American Highland vs. South American Lowland. *Q_ST_* values were considered significantly high if they were greater than two standard deviations from the mean *F_ST_*.

## 3 Results

### 3.1 Population Mean Reaction Norms

Reaction norms describe phenotypic trait values of genotypes (in this case, landrace populations) at different environments (common garden sites). Nonparallel reactions norms indicate that populations respond to environments differently, a pattern known as genotype-by-environment (*G* × *E*) interaction. When local populations have fitness trait values higher than the fitness trait values of non-local populations, resulting in crossed reaction norms, this is known as local adaptation.

A full report of the statistical significance of each contrast is provided in Table 2. Bonferroni correction (Bonferroni, 1936) is used to account for multiple comparisons (for tests of each trait within each contrast).

**Table 2:**
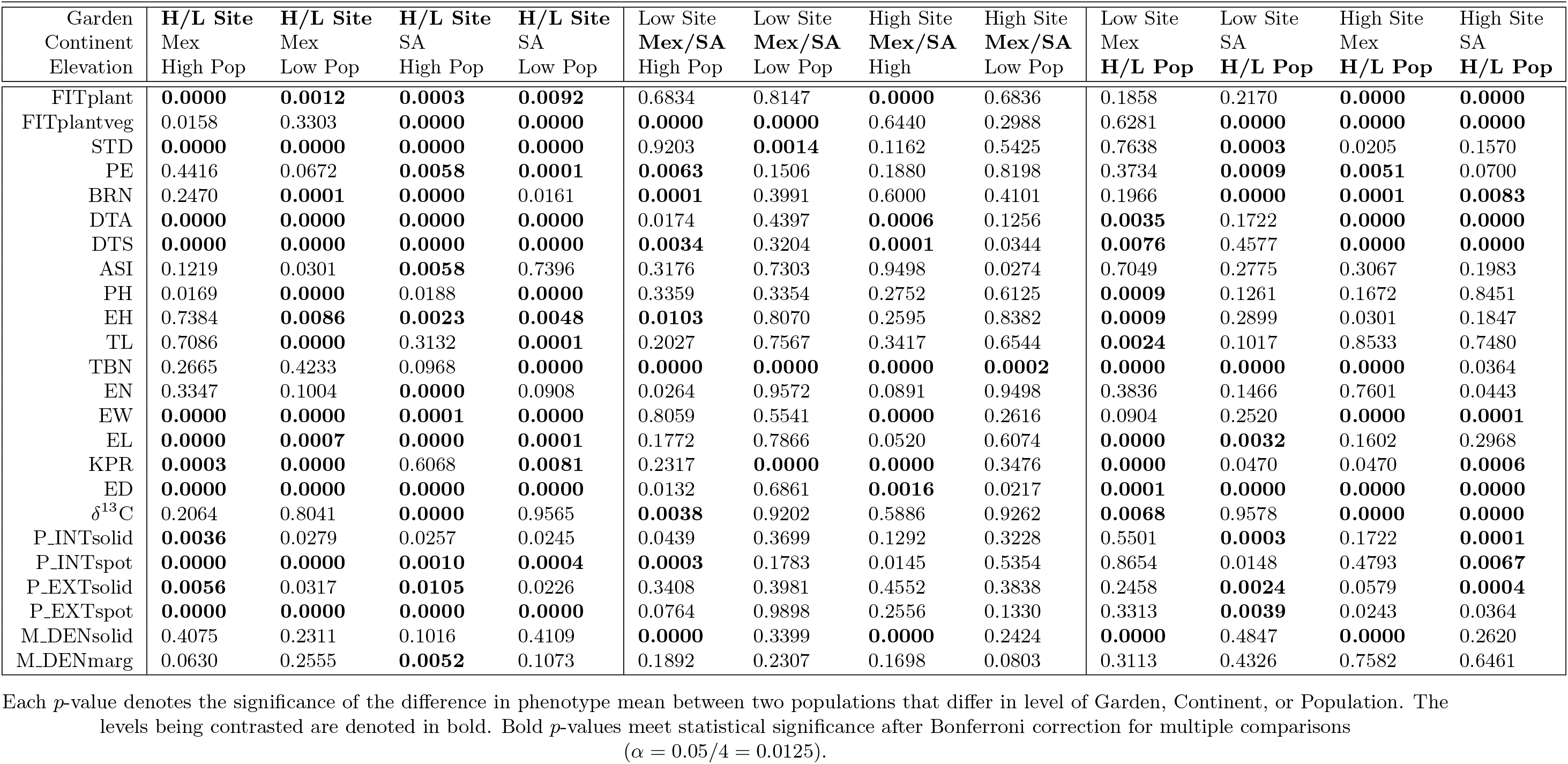
Reaction norm *p*-values

#### 3.1.1 Adjusted Fitness (FITplant, FITplantveg)

Both agronomic fitness (FITplant, Figure 2a) and vegetative fitness (FITplantveg, Figure 2b) showed strong patterns of home-site advantage. Though FITplant was highest in the High Site for all populations, both traits showed crossing reaction norms indicative of local adaptation.

**Figure 2:**
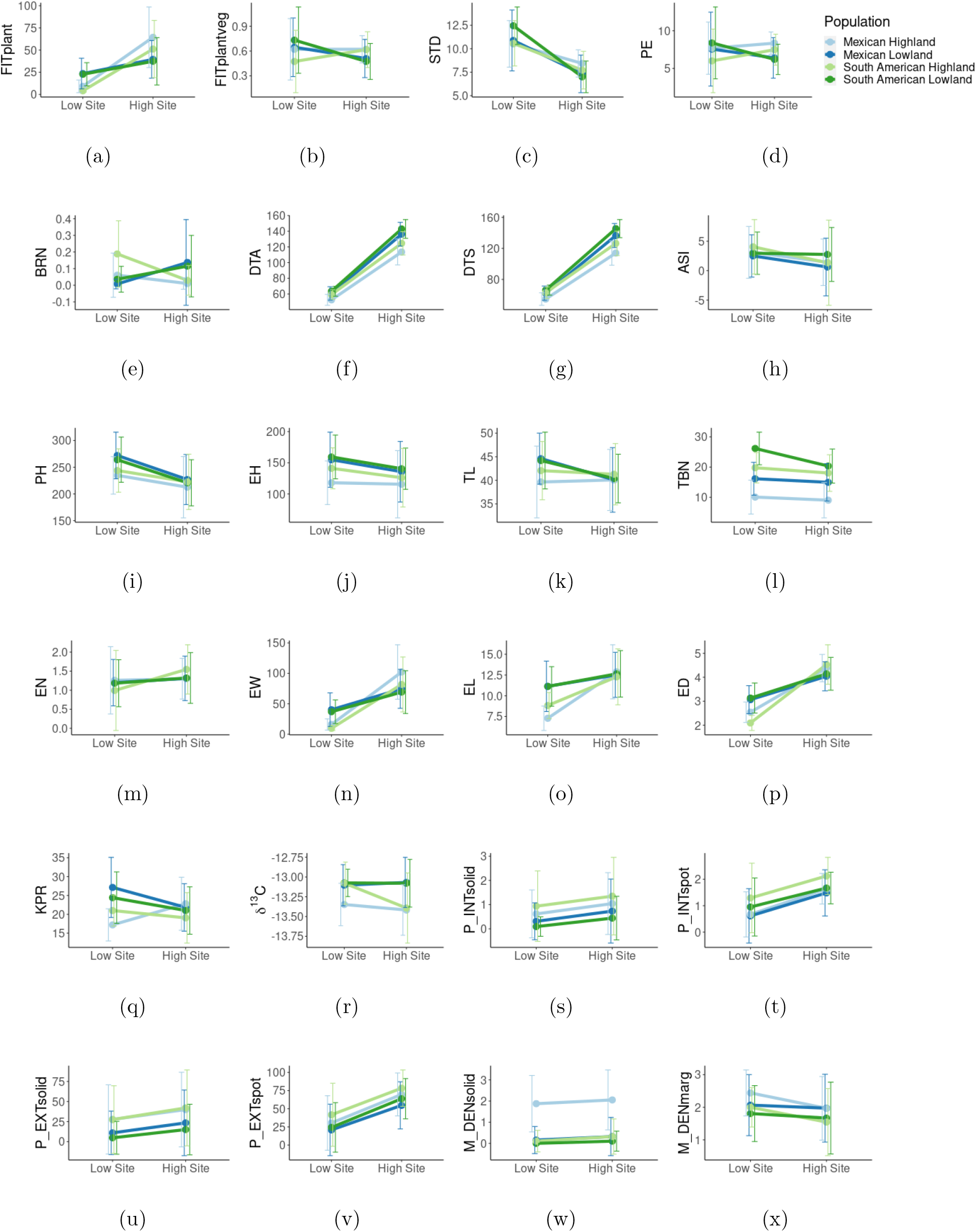
Reaction norms for all measured phenotypic traits.

Population values of FITplant in the Low Site were not significantly different from one another, though Low populations showed a modest advantage over High populations and Mex populations had an advantage over SA populations. In the High Site, these patterns crossed the significance threshold. In general, we would expect to see greater yield (and therefore greater FITplant) in the Low Site due to more tropical growth conditions, but these data showed the opposite trend. This was likely due to generally poor field conditions in the Low Site during the year that yield data were collected. For comparison, see ear weight (EW, Figure 2n).

FITplantveg values were largely reflective of ear-producing stand count (PE, Figure 2d). In the Low Site, Low populations had higher FITplantveg than High populations (though this difference was not statistically significant between Mex Low and Mex High), and Mex populations had higher fitness than SA populations. In the High Site, High populations had higher FITplantveg than Low populations, though Mex/SA differences were not significant.

#### 3.1.2 Row-Level Traits (STD, PE, BRN, DTA, DTS, ASI)

All four populations had significantly lower stand count (STD, Figure 2c) in the High Site than in the Low Site, though a weak pattern of home-site advantage emerged. Ear-producing stand count (PE, Figure 2d) showed stronger home-site advantage. SA populations crossed reaction norms, and all four populations had higher PE at their native elevation (though this trend was insignificant for Mex High). Notably, at the Low Site, PE values from Mex High and Mex Low nearly converged, whereas SA High and SA Low values diverged widely.

Barrenness (BRN, Figure 2e) is the percent of plants that survive but do not produce ears, whereas PE is the number of plants that germinate and survive to produce ears. BRN was negatively correlated with PE and therefore showed patterns similar but opposite to PE (lower BRN at native elevation).

Flowering time traits days to anthesis (DTA, Figure 2f) and days to silking (DTS, Figure 2g) showed that flowering took longer in the High Site. Though all populations showed similar patterns, South American populations took longer to flower than Mexican populations, and lowland populations took longer than highland populations. Anthesis/Silking Interval (ASI, Figure 2h) was generally lower in the High Site. The only significant ASI contrast was SA High between the Low and High Sites.

#### 3.1.3 Plant Size Traits (PH, EH, TL, TBN, EN)

Plant height (PH, Figure 2i) and ear height (EH, Figure 2j) were lower in the High Site than in the Low Site. Mex High was the only population that did not significantly vary between sites. Mex Low had higher PH and EH than Mex High in both sites. Lowland populations had much greater tassel length (TL, Figure 2k) in the Low Site than the High Site, but neither highland population varied substantially. Only SA Low varied between sites for tassel branch number (TBN, Figure 2l), but there was a strong genetic effect between populations in both sites. SA and Low populations had greater TBN than Mex and High populations. Ear number (EN, Figure 2m) was largely static between sites and between populations, except for SA High, which had a lower value in the Low Site and a higher value in the High Site.

#### 3.1.4 Yield Traits (EW, EL, ED, KPR)

Ear weight (EW, Figure 2n), ear length (EL, Figure 2o), and ear diameter (ED, Figure 2p) were all greater in the High Site than the Low Site. These depressed Low Site data trends may have been due in part to virus damage in the Low Site in 2016. EW and ED showed crossing reaction norms indicative of home-site advantage, and Mex populations had greater EW than SA populations from the same elevation. Low had greater EL than High populations in the Low Site, but values nearly converged in the High Site. Kernels per row (KPR, Figure 2q) did not exhibit the same depression at the Low Site as EW, EL, and ED. SA and Mex Low had lower KPR in the High Site, though Mex High had a strong opposite reaction norm.

#### 3.1.5 Water Use Efficiency (*δ*^13^C)

SA Low, SA High, and Mex High did not vary greatly for *δ*^13^C (*δ*^13^C, Figure 2r). Mex High had lower *δ*^13^C than both lowland populations in both sites. SA High showed a peculiar pattern of high *δ*^13^C in the Low Site, similar to both lowland populations, and low *δ*^13^C in the High Site, similar to Mex High.

#### 3.1.6 Anthocyanin Pigmentation and Macrohair Density (P_INTsolid, P_INTspot, P_EXTsolid, P_EXTspot, M_DENsolid, M_DENmarg)

Solid-pattern anthocyanin intensity (P_INTsolid, Figure 2s) and spot-pattern anthocyanin intensity (P_INTspot, Figure 2t) increased in the High Site relative to the Low Site. All four populations showed similar rates of increase between sites for both traits, but the increase in P_INTspot was statistically significant for all populations, and the increase in P_INTsolid was significant only for Mex High. In all cases, SA High had the highest intensity of both patterns of pigmentation, and in all cases, High populations had higher intensity than Low populations.

Solid-pattern anthocyanin extent (P_EXTsolid, Figure 2u) and spot-pattern anthocyanin extent (P_EXTspot, Figure 2v) increased in the High Site relative to the Low Site. The increase of P_EXTspot was significant for all populations, and the increase in P_EXTsolid was significant for High. In all cases, SA High had the highest pigmentation extent of both patterns of pigmentation, and in all cases, High had higher extents than Low.

Leaf sheath macrohair density (M_DENsolid, Figure 2w) and leaf sheath margin macrohair density (M_DENmarg, Figure 2x) demonstrated distinct patterns. None of the populations varied significantly in M_DENsolid between sites. Mex High had greater M_DENsolid than Mex Low and SA High in both sites, but otherwise, no significant differences were found. The only population that varied significantly in M_DENmarg between garden sites was SA High. No other significant differences were found for this trait.

### 3.2 Garden-Level Phenotypic Differences

Traits with low missing data between the three gardens (the High Site and both years of the Low Site) were used to perform Principal Components Analysis (Figure 3). The first two components distinguish individuals from the High Site from both plantings of the Low Site. The two years of the Low Site share a higher degree of feature space overlap than either shares with the High Site. High values of P_INTsolid, P _EXTsolid, DTA, and DTS characterize plants from the High Site. High values of several fitness-related traits and low values of M_DENsolid distinguish the Low Site 2017 from the Low Site 2016 and the High Site.

**Figure 3:**
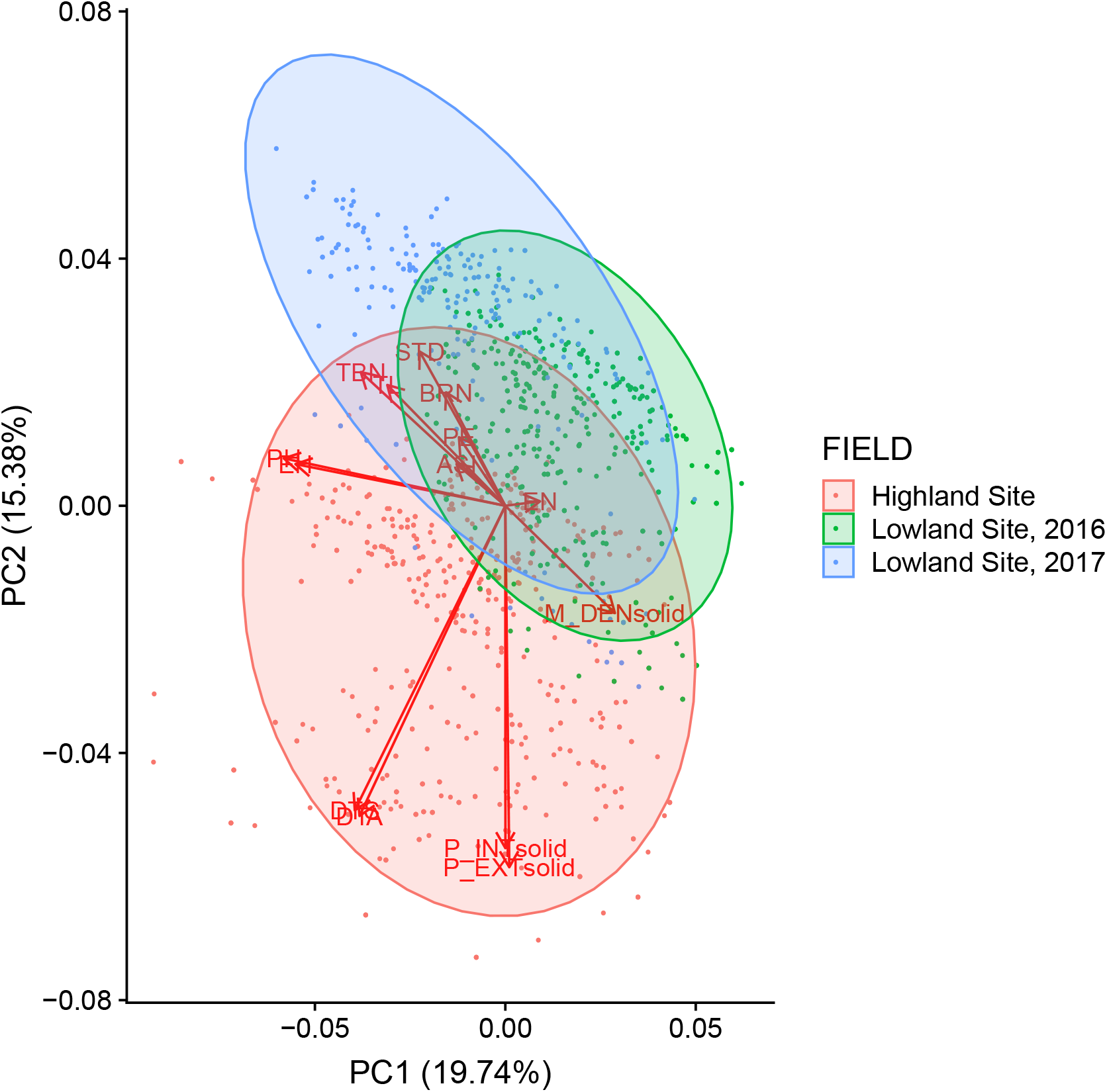
PCA of family and plant trait values between common garden plantings.

### 3.3 Pearson Correlation of Highland-Adaptive Traits

Pearson correlation values between fitness, pigment traits, and macrohair traits vary between all four populations and between both gardens. In all cases, P_INTsolid is positively correlated with P _EXTsolid (Figure 4), and P_INTspot is positively correlated with P_EXTspot (Figure 5). Likewise, in all cases, the strength of the correlation between P_INTsolid and P_EXTsolid is the same or greater in the High Site than in the Low Site (Figure 4). Conversely, the correlation between P_INTspot and P_EXTspot is weaker in the High Site than in the Low Site (Figure 5).

**Figure 4:**
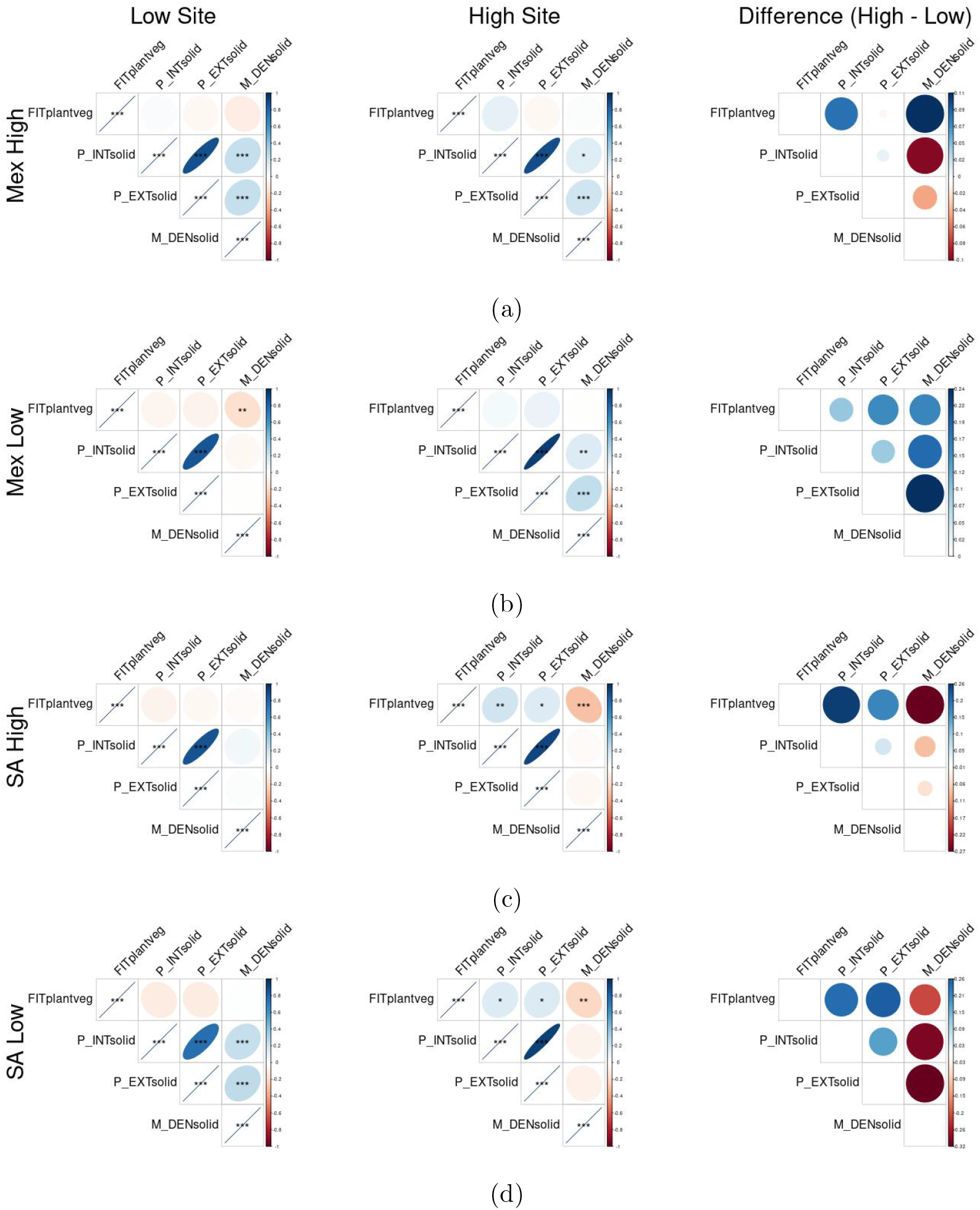
Pearson correlation between plant vegetative fitness, P_INTsolid, P_EXTsolid, and M_DENsolid in Mexican Highland (a), Mexican Lowland (b), South American Highland (c), and South American Lowland (d) populations. For each subfigure, panels 1 and 2 show correlations within the Low Site and the High Site. In panels 1 and 2, blue shapes indicate positive correlation, red shapes indicate negative correlation, color intensity and shape size indicate strength of correlation, and asterisks indicate statistical significance (*p*-value thresholds = 0.05, 0.01, 0.001). Panel 3 shows the between-garden difference in correlation value for each pairwise correlation (positive/blue values indicate more positive correlations in the highland site than in the lowland site).

**Figure 5:**
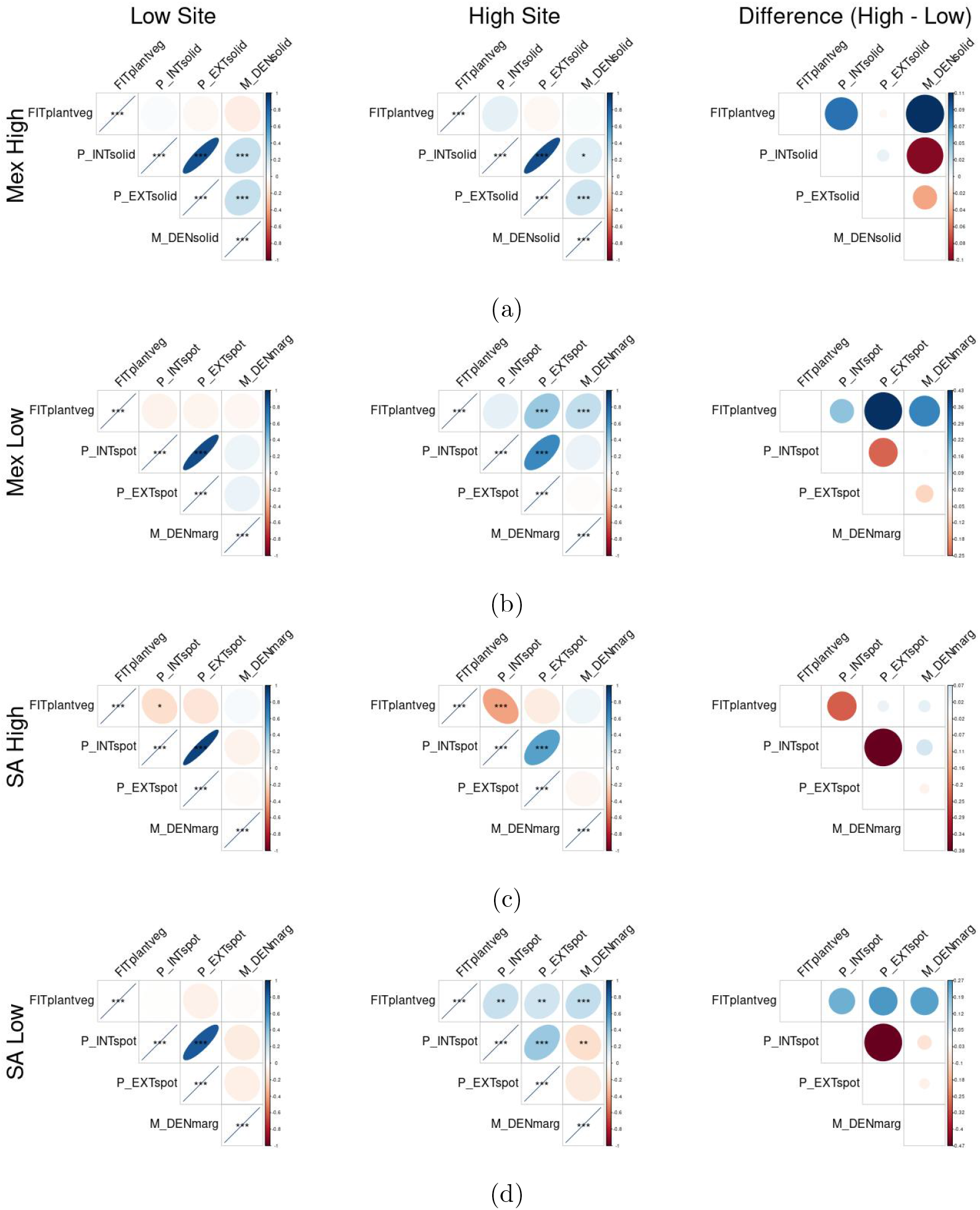
Pearson correlation between plant vegetative fitness, P_INTspot, P_EXTspot, and M_DENmarg in Mexican Highland (a), Mexican Lowland (b), South American Highland (c), and South American Lowland (d) populations. For each subfigure, panels 1 and 2 show correlations within the Low Site and the High Site. In panels 1 and 2, blue shapes indicate positive correlation, red shapes indicate negative correlation, color intensity and shape size indicate strength of correlation, and asterisks indicate statistical significance (*p*-value thresholds = 0.05, 0.01, 0.001). Panel 3 shows the between-garden difference in correlation value for each pairwise correlation (positive/blue values indicate more positive correlations in the highland site than in the lowland site).

#### 3.3.1 P_INTsolid, P_EXTsolid, and M_DENsolid

The correlations between M_DENsolid and either P_INTsolid or P_EXTsolid varies between populations and gardens, but is generally either weak (positive or negative) or strongly positive. In Mex High, these correlations are strongly positive in both gardens, though stronger in the Low Site. In Mex Low, these correlations are only strongly positive in the High Site. In SA Low, these correlations are only strong in the Low Site. In SA High, these correlations are weak in both gardens, but may be marginally positive in the Low Site and negative in the High Site.

In the Low Site, for all four populations, FITplantveg was either uncorrelated or negatively correlated with P_INTsolid, P_EXTsolid, and M_DENsolid, with the exception of a very weak positive correlation between FITplantveg and P_INTsolid within Mex High (Figure 4a, panel 1). Most of these negative correlations are weak, with the exception of a strong negative correlation between FITplantveg and M_DENsolid within Mex Low, Figure 4b, panel 1).

In contrast, in the High Site, for all four populations, FITplantveg was positively correlated with P_INTsolid, positively correlated with P_EXTsolid (except for Mex Low), and either uncorrelated or negatively correlated with M_DENsolid. However, the only significant correlations listed above were found in the SA High population.

Panel 3 in each subfigure of Figure 4 demonstrates difference in correlation between the Low and High sites. For all populations, the correlation between FITplantveg and P_INTsolid is stronger in the High Site than in the Low Site. The same is true of P_EXTsolid, except for Mex High, in which there is no change in correlation between garden sites. On the other hand, Mexican populations show an increase in correlation between FITplantveg and M_DENsolid in the High Site relative to the Low Site, and South American populations show the opposite trend. Also, all four populations except for Mex Low show weaker correlation between M_DENsolid and either P_INTsolid or P_EXTsolid in the High Site relative to the Low Site (though the Mex High correlation coefficients are still both significantly positive in both sites).

#### 3.3.2 PINTspot, P_EXTspot, and MDENmarg

In both gardens, M_DENmarg is negatively correlated with P_INTspot and P_EXTspot in South American populations, and weakly or positively correlated in Mexican populations. In the Low Site, P_INTspot, P_EXTspot, and M_DENmarg are weakly or negatively correlated with FITplantveg in all populations. This correlation becomes positive for the Low populations in the High Site, but patterns in the High populations are mixed.

### 3.4 Population Genetic Relatedness

Genetic similarity between genotyped individuals based on SNP data is estimated with Principal Components Analysis (Figure 6a). Component 1 (24.7%) primarily separates Mexican from South American populations, and Component 2 (14.4%) primarily separates highland from lowland populations.

**Figure 6:**
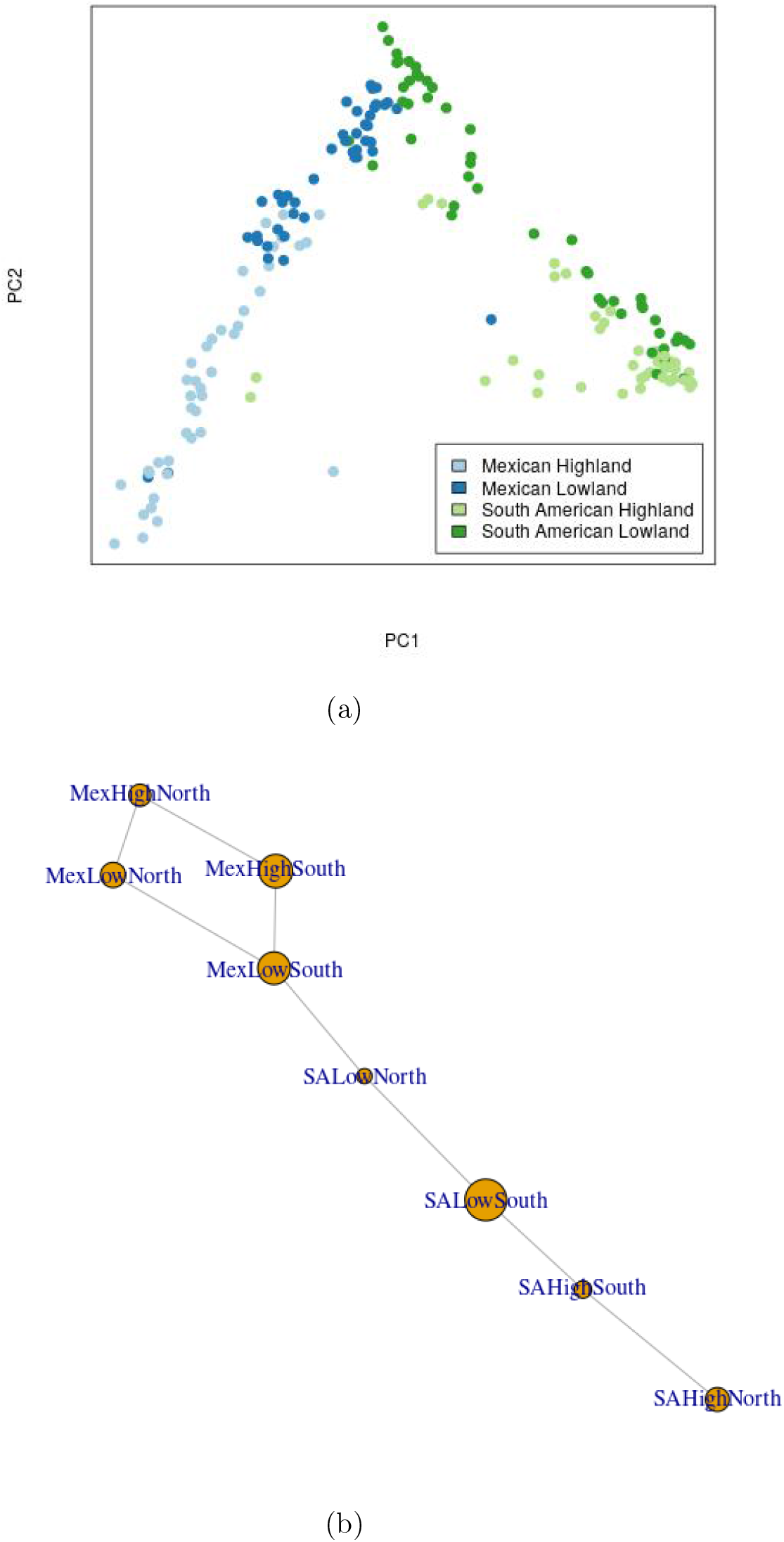
Population genetic structure among maize landrace populations. (6a) Principal Components Analysis of SNPs (Component 1 = 24.7%, Component 2 = 14.4%). (6b) Population graph of eight continent/elevation/latitude subpopulations derived from four continent/elevation populations. Node size reflects degree of genetic variability within a subpopulation, and edge length reflects among-subpopulation genetic variance.

The population graph (Figure 6b) likewise separates the eight subpopulations into Mexican and South American groups. Greater edge saturation amongst Mexican subpopulations illustrates greater genetic similarity amongst Mexican subpopulations than amongst South American sub-populations. Differential node sizes reflect the degree of genetic variability within subpopulations. The subpopulations with the greatest genetic variability are MexHighSouth, MexLowSouth, and SALowSouth.

### 3.5 *Q_ST_/F_ST_* Comparison

*Q_ST_* values for quantitative traits were plotted against the distribution of *F_ST_* values (Figure 7). Four sets of comparisons were carried out; High vs. Low (Figure 7a), Mex vs. SA (Figure 7b), Mex High vs. Mex Low (Figure 7c), and SA High vs. SA Low (Figure 7d). *Q_ST_* values more than two standard deviations above the mean *F_ST_* were considered significantly high. In all three elevational comparisons, PH, EH, DTS, DTA, and *δ*^13^C had significantly high *Q_ST_. Q_ST_* values for TL, TBN, P_INTsolid, M_DENsolid also meet this threshold of significance in one or two of the elevational contrasts. In the Mex vs. SA contrast, only three traits (TBN, M_DENsolid, and DTS) have significantly high *Q_ST_* values.

**Figure 7:**
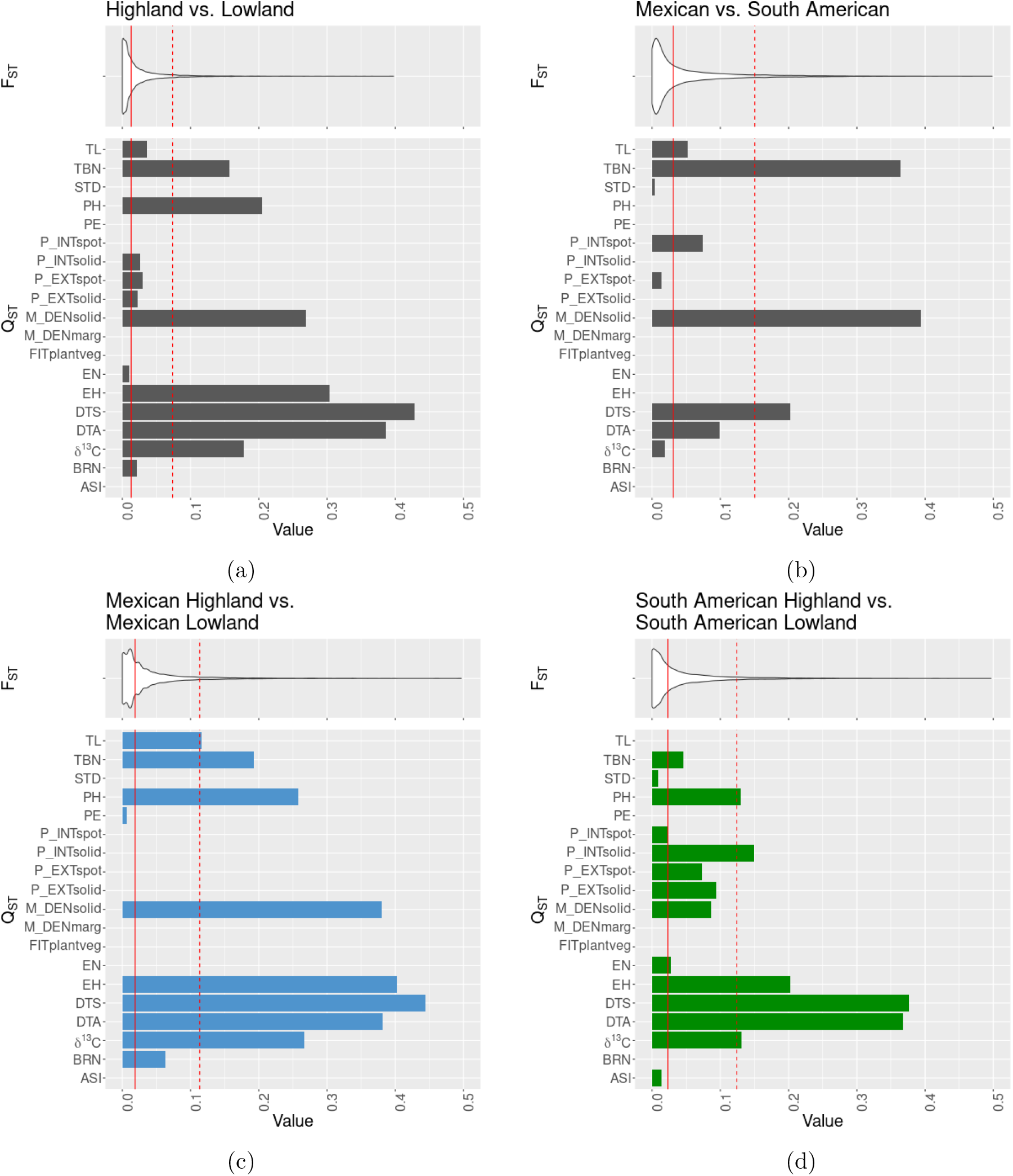
*F_ST_* and *Q_ST_* values between four sets of populations. Solid red lines indicate mean *F_ST_* and dashed red lines indicate two standard deviations from the mean. (7a) Highland vs. Lowland. (7b) Mexican vs. South American. (7c) Mexican Highland vs. Mexican Lowland. (7d) South American Highland vs. South American Lowland.

## 4 Discussion

### 4.1 Local Adaptation and Plasticity

Landraces may respond to environmental changes (including climate change and dispersal to new environments) in up to four ways: plasticity, evolution, gene flow, or extinction (Mercer and Perales, 2010). The failure of an organism to plastically adapt to all available environments promotes the evolution of adaptations to a particular environment at the expense of others, a compromise known as an adaptive trade-off. When a population evolves traits that give it a home-site advantage over non-native populations, that population exhibits local adaptation (Kawecki and Ebert, 2004).

All four of our elevational/continental landrace populations differ in fitness component values between our highland and lowland Mexican field sites. We observe that populations exhibit reciprocal home-site advantage in several ways. Populations grown at sites near their native elevation have higher agronomic and vegetative fitness, stand count, ear-producing stand count, ear weight, ear diameter, and lower barrenness than populations foreign to that site’s elevation, as indicated by crossing reaction norms between populations from the same continent. Other traits show evidence of home-site advantage for populations from one continent, but not the other, indicating that highland and lowland populations from different continents have different adaptive strategies.

In several cases, populations also fit the “Home vs. Away” model of local adaptation, in which a population has greater fitness in the site corresponding to its native home environment than in the away site, regardless of the fitness of other populations. The Mexican Highland and Lowland populations demonstrate this pattern most clearly with ear-producing stand count (Figure 2d). Though their reaction norms do not cross, both populations have higher fitness in their home sites. We might consider that, when populations meet the requirements for both models of local adaptation, there is a particularly strong case for local adaptation.

Several traits showed strong environmental effects but minimal *G* × *E*. All populations respond similarly to site effects for several traits, including days to anthesis, days to silking, plant height, and to lesser extents, spot-pattern pigment intensity and extent. These results are in alignment with expectations of depressed maize plant height and prolonged maturation process due to highland conditions (Mercer and Perales, 2018; Hufford et al., 2013).

We note that our reciprocal transplant design is not fully reciprocal in that common garden sites in South American locales were not utilized. Though we may expect to see South American populations exhibiting higher fitness than Mexican populations in such locales, this is currently speculative.

### 4.2 Highland Adaptation Traits

#### 4.2.1 Anthocyanin Pigmentation and Macrohair Density

Leaf sheath anthocyanin pigmentation and pilosity have long been reported to help plants acquire and retain heat in cold environments (Doebley, 1984; Schuepp, 1993; Lauter et al., 2004). Anthocyanin pigmentation is plastically up-regulated in response to increased light exposure (Vanderauwera et al., 2005) and cold temperatures (Christie et al., 1994; Hufford et al., 2013). We find that the intensity and extent of anthocyanin pigmentation on leaf sheaths is elevated in the highland garden site. In general, highland populations have greater overall pigmentation intensity and extent, though all populations demonstrate similar plastic effects in response to environment. *Q_ST_* of solid-pattern anthocyanin intensity is significantly high between South American Highland and South American Lowland, signifying differential directional selection for this trait between these two populations (Merilä and Crnokrak, 2001), but Mexican Highland and Mexican Lowland have low *Q_ST_* for this trait. Also, a difference in pattern emerges between Lowland populations, wherein Mexican Lowland has greater intensity and extent of solid-pattern anthocyanin, and South American Lowland has greater intensity and extent of anthocyanin spots. The correlations between both patterns of anthocyanin and fitness appear to become more positive with increasing elevation, though solid anthocyanin pigmentation has a somewhat more positive correlation with fitness than does anthocyanin spots.

Macrohair density is a plastic trait, associated with survival in cold temperatures (Hufford et al., 2013) and with maize grain yield in cold environments (Kaur et al., 1985). Unexpectedly, leaf sheath macrohair density was largely non-plastic to the environmental variation present in this study. Leaf sheath macrohair density is much greater in Mexican Highland maize than in the other populations, and this difference is greater than expected given neutral genetic loci. The introgression of macrohair density QTLs from *mexicana* exlusively into Highland Mexican maize (Lauter et al., 2004), followed by selection for that phenotype in the highland environment, would account for this pattern.

The fitness correlations with macrohair density are different in Mexican and South American maize landraces. The correlation between plant fitness and leaf sheath macrohair density becomes stronger at the highland site for Mexican landrace populations, but becomes more negative for South American landrace populations. This may suggest that leaf sheath macrohairs are adaptive for Mexican landraces in the Mexican Central Plateau highlands, but are not adaptive for South American landraces in this environment. The marginally higher density of sheath margin macrohairs among Mexican populations relative to South American populations adds support to this conclusion. Because leaf sheath macrohair density is (weakly) negatively correlated with anthocyanin pigmentation in the highland site for South American populations, and positively correlated with anthocyanin pigmentation intensity for Mexican populations in the highland site, this macrohair/fitness correlation may simply be a reflection of the fitness consequences of anthocyanin pigmentation. On the other hand, macrohair’s negative correlation with fitness is stronger than its negative correlation with anthocyanin intensity, suggesting some relationship between macrohair density and fitness beyond conflation with anthocyanin pigment. Furthermore, as variation in macrohair density is mainly captured by landrace population, this trait’s correlation with other traits will capture other population-level patterns in trait/trait correlations.

The only population in which (solid) anthocyanin pigmentation and leaf sheath macrohair density are strongly correlated regardless of environment is Mexican Highland. Whereas the other three populations show weaker or environmentally conditional correlation between anthocyanin and leaf sheath macrohair density, Mexican Highland maize shows the expression of these traits to be linked. Linkage mapping experiments would reveal whether the QTLs for these traits are truly under linkage disequilibrium, perhaps within the well-known inversion polymorphism, *inv4m* (Hufford et al., 2013; Pyhäjärvi et al., 2013), that overlaps with QTL for these traits (Lauter et al., 2004).

#### 4.2.2 Flowering Time/Plant Maturation

Flowering time is a complex, multigenic trait that plays a crucial role in elevation adaptation (Buckler et al., 2009; Navarro et al., 2017; Wang et al., 2020). Fast flowering time is a critical component of adaptation to cold highland conditions, as plants must complete their life cycle in a narrower window of hospitable weather. In accordance with these expectations, highland populations matured more quickly than lowland populations, and this difference was more pronounced in the highland site. At the same time, maize plants from all four populations had longer flowering time in the highland site, due to the slower accumulation of growing degree days. Strong signals of *Q_ST_* > *F_ST_* support strong divergent selection between highland and lowland population for flowering time.

Positive values of ASI indicate pollen release before silks are developed and receptive, which can lead to incomplete pollination and reduced yield. Positive values of ASI negatively correlate with yield (Mercer and Perales, 2018), but low or slightly negative values of ASI are likely less detrimental, as silks can remain receptive for several days, and a single plant that is shedding pollen early can pollinate many plants. For this reason, high values of ASI are generally regarded as an indicator of stress (Mercer et al., 2014). All four populations showed slightly higher ASI in the lowland site (though only South American Highland varied significantly). ASI reaction norms for Mexican populations are roughly parallel, while the South American reaction norms cross. This may be because ASI is more associated with local adaptation strategies of South American populations, or it may be that ASI is sensitive to the compounding stress of trans-elevational and trans-continental transplantation (Mittler, 2006). Both South American populations have ASI values resembling Mexican populations of the same elevation when grown at their native elevation, and then deviate more strongly when grown at the alternative elevation.

#### 4.2.3 Plant Morphology and Architecture

In maize, height is a polygenic trait with broad-ranging fitness consequences (Lin et al., 1995). Lowland populations are taller than Highland populations, and this difference is greater than expected given neutral genetic markers. Maize plant height is both highly heritable (Peiffer et al., 2014) and highly plastic to environment: All populations were shorter when grown in the highland site, reflecting the environmental effect of the shorter growing season at the highland site.

Optimal tassel size requires a tassel small enough for minimization of shading effects on the upper leaves yet large enough for sufficient pollen production (Mickelson et al., 2002), though the adaptive significance of tassel morphology is not well-known. Our *Q_ST_/F_ST_* comparisons reveal that Lowland populations have more branches than Highland populations (as observed by Eagles and Lothrop (1994)) and that South American populations have more branches than Mexican populations. Though tassel lengths of Highland populations were largely non-plastic, Lowland populations experienced a significant reduction in tassel length when transplanted to the lowland garden. Only the South American Lowland population was plastic between garden sites.

#### 4.2.4 Water Use Efficiency and *δ*^13^C

In C_4_ plants like maize, there is a negative correlation between WUE and *δ*^13^C (Ellsworth and Cousins, 2016). Individuals with higher/less negative *δ*^13^C scores have higher ratios of ^13^C:^12^C, meaning that they discriminate less effectively against ^13^C. Though the precise mechanism underlying this relationship is unclear, Avramova et al. (2019) found a region on Chromosome 7 which influences *δ*^13^C, WUE, and sensitivity to drought through reduced abscisic acid and modified stomatal behavior. Because precipitation decreases with increasing elevation in Mexico and South America, higher WUE may play a role in highland adaptation.

Both Lowland populations show consistently high *δ*^13^C, indicating low WUE. The Mexican Highland population had consistently lower *δ*^13^C at both sites, indicating higher WUE. This finding is in accord with other published studies that detail the various drought-adapted landraces of the Mexican highlands (Eagles and Lothrop, 1994; Hayano-Kanashiro et al., 2009). In both Mexican Highland/Mexican Lowland and South American Highland/South American Lowland comparisons, *Q_ST_* > *F_ST_*, indicating differential selection on WUE between highland and lowland populations on both continents. Only the South American Highland population differed for *δ*^13^C significantly between sites. South American Highland maize, like Mexican Highland maize, had high WUE in the highland site, but WUE dropped significantly in the lowland site. This unexpected drop in WUE seen in South American Highland maize may be the result of accumulated stress from being outside its native elevation and continent, though similar extreme drops in values of other fitness-relevant traits in the South American Highland population are not observed.

### 4.3 Population Structure

For our provisional formulation of four maize landrace populations divided by continent and elevation, populations are more genetically similar to the corresponding population from the same continent. This is demonstrated by the genetic PCA, in which PC1 most clearly distinguishes Mexican from South American landraces. The patterns observed in this PCA are congruent with those found by Kistler et al. (2018) in a diverse array of maize landrace and teosinte accessions from across the Americas, though their study divided accessions into population groups with a model-based clustering algorithm.

Dissecting our four landrace populations latitudinally, the population graph clusters the four Mexican subpopulations into one group and the four South American subpopulations into another, with a single edge between the two groups. This cluster pattern conforms to expectations, given the geographic proximity of these subpopulations and historical range expansion of maize from the center of domestication in southern lowland Mexico (MexLowSouth) to the north (MexLowNorth) and to the south (all South American subpopulations) and to higher elevations (MexHighSouth). Within either continent, subpopulations are most related to the subpopulation of the same elevation, rather than latitude, demonstrating that elevation structures genetic covariance more strongly than latitude. The two southern Mexican subpopulations have slightly higher within-subpopulation genetic variance than the two northern Mexican subpopulations, which reflects their closer proximity to the center of domestication and diversity (Wang et al., 2017). On the other hand, this pattern is not shown within the South American subpopulations; SALowSouth has greater genetic variance than does SALowNorth. This result is in alignment with the hypothesis that the southwestern Amazon (present-day Bolivia, Peru, and southwestern Brazil), which roughly corresponds to the geographical region of SALowSouth, was a plant domestication and production hotspot (Watling et al., 2018) and secondary improvement center for maize (Kistler et al., 2018). According to this theory, this region was the destination of the first of two waves of maize dispersal to South America, and from this region, maize dispersed across the rest of the continent, including to the northern South American lowlands. Given this hypothesis, we would expect MexLowSouth to have a stronger edge to SALowSouth than to SALowNorth. There are two possible reasons why we do not find this to be the case. First, this genetic covariance may be preserved from the original southward dispersal of maize from the center of domestication to the secondary improvement center through the northern lowlands. The second and more likely scenario is that this stronger covariance reflects gene flow from Mesoamerica (MexLowSouth) to South America (SALowNorth) across Central America sometime after the first wave of dispersal of maize into South America, either at low consistent levels or as part of a second wave of dispersal (Kistler et al., 2018, 2020).

Though populations are more genetically similar to the corresponding population from the same continent, they are phenotypically more similar to the populations from the same elevation. While genetic population structure is largely shaped by demographic effects of drift during dispersal, phenotypes and phenotypic plasticity show evidence of being shaped by elevational adaptation. This is apparent for the majority of traits’ reaction norms between common garden sites, as well as the generally higher *Q_ST_* between highland and lowland populations than that between Mexican and South American populations.

### 4.4 Asymmetrical Patterns of Local Adaptation

Mercer et al. (2008) found that highland populations suffer a greater reduction in fitness in lowland conditions than lowland populations do in highland conditions. They describe this pattern as asymmetrical local adaptation. Our data do not fully replicate this finding. Our agronomic fitness data approach this pattern, with relatively stable lowland fitness and more variable highland fitness, but vegetative fitness shows an opposite asymmetry with more variable lowland populations and more stable highland populations. As Mercer and colleagues focused on agronomic fitness, these results are in alignment. Any asymmetry of local adaptation found here may be sensitive to yearly fluctuations in *G* × *E* interactions at a site (Mercer and Perales, 2018). Further studies would be required (and are recommended) to see whether patterns of asymmetry break down or are retained over time.

### 4.5 Selective Forces in Maize Evolution

Agroecosystems exert multiple and at times conflicting selective pressures on maize populations. Fitness is defined as (or approximated by, Savolainen et al. (2013)) an organism’s ability to survive and reproduce successfully in a particular environment. Fit maize plants must survive the myriad forces at work in the field (due to climate, elevation, soil type and quality, pest and weed pressure, as well as farmer-mediated modifications to the land, such as tilling, irrigation, fertilizer, and crop rotation) to germinate, mature, develop numerous healthy seeds, and resist post-harvest spoilage and loss. Furthermore, fit maize plants must also satisfy the desires of farmers to such a degree that the farmers will be convinced to replant the seed line in subsequent seasons. In fact, farmers more commonly report consciously selecting for culinary traits than for yield or environmental adaptations (Bellon et al., 2003). While maize populations continually evolve in response to competing selective pressures, agronomic practices and consumption patterns also evolve to maximize yield, minimize required inputs, and produce seed with desired grain type.

Though highland-adapted landraces in Mexico and South America share phenotypic similarities, their adaptive strategies are not identical. This is evinced by highly divergent reaction norms between Mexico and South America for a few traits, notably *δ*^13^C. Differences in highland adaptation between Mexican and South American maize may be due to drift incurred during the dispersal of landraces into and across South America, the unique selective challenges imparted by specific local highland regions, or likely a combination of both.

The diversity and complexity of selective forces at work in the maize landrace agroecosystem may impede detection of patterns of adaptation to abiotic clines like elevation, which may explain why the common garden experiment by Orozco-Ramírez et al. (2014) failed to identify environmental adaptation as a leading factor in landrace distribution, and why the analyses of Dyer and López-Feldman (2013) found that altitude did not cleanly explain seed management practices. The clear patterns of adaptation to elevation found in this reciprocal transplant experiment are perhaps more striking when considering the complicating and significant force of anthropogenic (or “artificial”) selection.

Additionally, the common garden sites were maintained at similar modern agronomic conditions (irrigation and pesticide/insecticide/fungicide inputs). This is in contrast to the diverse traditional agronomic practices utilized at smallholder farms across Mexico in which landrace diversity is maintained. This disparity between native habitat conditions and common garden conditions may have further reduced the observable signal of local adaptation to mitigated selection pressures (e.g., drought and biotic pressures), but others environmental pressures (temperature, ultraviolet solar radiation, atmospheric pressure, etc.) are less likely to have been affected.

## 5 Conclusions

These results demonstrate that maize landraces from across the Americas are locally adapted to elevation. Landraces adapted to diverse environmental conditions are an invaluable resource for breeding efforts that rely on fewer costly and ecologically harmful inputs (Dwivedi et al., 2016). The myriad forces that influence the *in situ* conservation status of landraces are complex and dynamic, though locally adapted and evolving populations are more resilient and less likely to be supplanted by modern varieties (Perales et al., 2003). The importance of landraces as an agronomic resource is likely to increase due to growing global food demands, the proliferation of modern inbred lines, and the effects of global climate change, which will likely alter the conditions of many corn-producing regions substantially (Bassu et al., 2014; Xu et al., 2016).

## Data Archiving Statement

Data for this study are available at: *to be completed after manuscript is accepted for publication*.

## References

Jonás A Aguirre-Liguori, Santiago Ramírez-Barahona, Peter Tiffin, and Luis E Eguiarte. Climate change is predicted to disrupt patterns of local adaptation in wild and cultivated maize. Proceedings of the Royal Society B, 286(1906):20190486, 2019.

María Clara Arteaga, Alejandra Moreno-Letelier, Alicia Mastretta-Yanes, Alejandra Vázquez-Lobo, Alejandra Breña-Ochoa, Andrés Moreno-Estrada, Luis E Eguiarte, and Daniel Piñero. Genomic variation in recently collected maize landraces from mexico. Genomics data, 7:38–45, 2016.

Viktoriya Avramova, Adel Meziane, Eva Bauer, Sonja Blankenagel, Stella Eggels, Sebastian Gresset, Erwin Grill, Claudiu Niculaes, Milena Ouzunova, Brigitte Poppenberger, et al. Carbon isotope composition, water use efficiency, and drought sensitivity are controlled by a common genomic segment in maize. Theoretical and Applied Genetics, 132(1):53–63, 2019.

Simona Bassu, Nadine Brisson, Jean-Louis Durand, Kenneth Boote, Jon Lizaso, James W Jones, Cynthia Rosenzweig, Alex C Ruane, Myriam Adam, Christian Baron, et al. How do various maize crop models vary in their responses to climate change factors? Global change biology, 20 (7):2301–2320, 2014.

Douglas Bates, Martin Mächler, Ben Bolker, and Steve Walker. Fitting linear mixed-effects models using lme4. arXiv preprint arXiv:1406.5823, 2014a.

Douglas Bates, Martin Maechler, Ben Bolker, and Steven Walker. lme4: Linear mixed-effects models using eigen and s4. r package version 1.1–7, 2014b.

Mauricio R Bellon, Julien Berthaud, Melinda Smale, José Alfonso Aguirre, Suketoshi Taba, Flavio Aragón, Jaime Díaz, and Humberto Castro. Participatory landrace selection for on-farm conservation: An example from the central valleys of oaxaca, mexico. Genetic Resources and Crop Evolution, 50(4):401–416, 2003.

Mauricio R Bellon, Alicia Mastretta-Yanes, Alejandro Ponce-Mendoza, Daniel Ortiz-Santamaría, Oswaldo Oliveros-Galindo, Hugo Perales, Francisca Acevedo, and José Sarukhán. Evolutionary and food supply implications of ongoing maize domestication by mexican campesinos. Proc. R. Soc. B, 285(1885):20181049, 2018.

Carlo Bonferroni. Teoria statistica delle classi e calcolo delle probabilita. Pubblicazioni del R Istituto Superiore di Scienze Economiche e Commericiali di Firenze, 8:3–62, 1936.

Mariana Bracco, Veronica Viviana Lia, J CÁMARA Hernáindez, Lidia Poggio, and Alexandra Marina Gottlieb. Genetic diversity of maize landraces from lowland and highland agro-ecosystems of southern south america: implications for the conservation of native resources. Annals of Applied Biology, 160(3):308–321, 2012.

Stephen B Brush. Man’s use of an andean ecosystem. Human Ecology, 4(2):147–166, 1976.

Edward S Buckler, James B Holland, Peter J Bradbury, Charlotte B Acharya, Patrick J Brown, Chris Browne, Elhan Ersoz, Sherry Flint-Garcia, Arturo Garcia, Jeffrey C Glaubitz, et al. The genetic architecture of maize flowering time. Science, 325(5941):714–718, 2009.

Mark B Bush, R Piperno Dolores, and Paul A Colinvaux. A 6,000 year history of amazonian maize cultivation. Nature, 340(6231):303, 1989.

Kridsadakorn Chaichoompu, Fentaw Abegaz, Sissades Tongsima, Philip James Shaw, Anavaj Sakuntabhai, Luisa Pereira, and Kristel Van Steen. KRIS: Keen and Reliable Interface Sub-routines for Bioinformatic Analysis, 2018. URL https://CRAN.R-project.org/package=KRIS. R package version 1.1.1.

Linda Chalker-Scott. Environmental significance of anthocyanins in plant stress responses. Photochemistry and photobiology, 70(1):1–9, 1999.

Peter J Christie, Mark R Alfenito, and Virginia Walbot. Impact of low-temperature stress on general phenylpropanoid and anthocyanin pathways: enhancement of transcript abundance and anthocyanin pigmentation in maize seedlings. Planta, 194(4):541–549, 1994.

Jens Clausen, David D Keck, William M Hiesey, et al. Experimental studies on the nature of species. i. effect of varied environments on western north american plants. Experimental studies on the nature of species. I. Effect of varied environments on western North American plants., (520), 1940.

David A Cleveland and Daniela Soleri. Extending darwins analogy: bridging differences in concepts of selection between farmers, biologists, and plant breeders. Economic botany, 61(2):121, 2007.

John F Doebley. Maize introgression into teosinte-a reappraisal. Annals of the Missouri Botanical Garden, pages 1100–1113, 1984.

Christian Dumas and H Lloyd Mogensen. Gametes and fertilization: Maize as a model system for experimental embryogenesis in flowering plants. The Plant Cell, 5(10):1337, 1993.

Sangam L Dwivedi, Salvatore Ceccarelli, Matthew W Blair, Hari D Upadhyaya, Ashok K Are, and Rodomiro Ortiz. Landrace germplasm for improving yield and abiotic stress adaptation. Trends in plant science, 21(1):31–42, 2016.

George A Dyer and Alejandro López-Feldman. Inexplicable or simply unexplained? the management of maize seed in mexico. PLoS One, 8(6):e68320, 2013.

RJ Dyer. popgraph: This is an r package that constructs and manipulates population graphs. R package version, 1, 2014.

Rodney J Dyer and John D Nason. Population graphs: the graph theoretic shape of genetic structure. Molecular ecology, 13(7):1713–1727, 2004.

HA Eagles and JE Lothrop. Highland maize from central mexicoits origin, characteristics, and use in breeding programs. Crop Science, 34(1):11–19, 1994.

Patrick Z Ellsworth and Asaph B Cousins. Carbon isotopes and water use efficiency in c4 plants. Current Opinion in Plant Biology, 31:155–161, 2016.

Nina Fedoroff. How jumping genes were discovered. Nature Structural & Molecular Biology, 8(4): 300, 2001.

Rocio Fernandez Suarez, Luis A Morales Chavez, and Amanda Galvez Mariscal. Importance of mexican maize landraces in the national diet. an essential review. Revista Fitotecnia Mexicana, 36:275–283, 2013.

Dylan J Fraser, Laura K Weir, Louis Bernatchez, Michael Møller Hansen, and Eric B Taylor. Extent and scale of local adaptation in salmonid fishes: review and meta-analysis. Heredity, 106(3):404, 2011.

Joseph L Gage, Diego Jarquin, Cinta Romay, Aaron Lorenz, Edward S Buckler, Shawn Kaeppler, Naser Alkhalifah, Martin Bohn, Darwin A Campbell, Jode Edwards, et al. The effect of artificial selection on phenotypic plasticity in maize. Nature communications, 8(1):1348, 2017.

Daniel J Gates, Dan Runcie, Garret M Janzen, Alberto Romero Navarro, Martha Willcox, Kai Sonder, Samantha Snodgrass, Fausto Rodríguez-Zapata, Ruairidh JH Sawers, Rubén Rellán-Álvarez, et al. Single-gene resolution of locally adaptive genetic variation in mexican maize. bioRxiv, page 706739, 2019.

Alexis L Gibson, Erin K Espeland, Viktoria Wagner, and Cara R Nelson. Can local adaptation research in plants inform selection of native plant materials? an analysis of experimental methodologies. Evolutionary Applications, 9(10):1219–1228, 2016.

Alexander Grobman, Duccio Bonavia, Tom D Dillehay, Dolores R Piperno, José Iriarte, and Irene Holst. Preceramic maize from paredones and huaca prieta, peru. Proceedings of the National Academy of Sciences, 109(5):1755–1759, 2012.

Bernd Gruber, Peter J Unmack, Oliver F Berry, and Arthur Georges. dartr: an r package to facilitate analysis of snp data generated from reduced representation genome sequencing. Molecular ecology resources, 18(3):691–699, 2018.

Jack R Harlan. Our vanishing genetic resources. Science, 188(4188):618–621, 1975.

Corina Hayano-Kanashiro, Carlos Calderión-Váizquez, Enrique Ibarra-Laclette, Luis Herrera-Estrella, and June Simpson. Analysis of gene expression and physiological responses in three mexican maize landraces under drought stress and recovery irrigation. PLoS one, 4(10):e7531, 2009.

Matthew B Hufford, Enrique Martínez-Meyer, Brandon S Gaut, Luis E Eguiarte, and Maud I Tenaillon. Inferences from the historical distribution of wild and domesticated maize provide ecological and evolutionary insight. PLoS One, 7(11):e47659, 2012.

Matthew B Hufford, Pesach Lubinksy, Tanja Pyhäjärvi, Michael T Devengenzo, Norman C Ellstrand, and Jeffrey Ross-Ibarra. The genomic signature of crop-wild introgression in maize. PLoS Genetics, 9(5):e1003477, 2013.

Garrett M Janzen, Li Wang, and Matthew B Hufford. The extent of adaptive wild introgression in crops. New Phytologist, 2018.

GP Kaur, M Dhillon, and BS Dhillon. Agronomic and anatomical characters in relation to cold-tolerance and grain-yield in maize composite. Proceedings Indian National Sciences Academy and Biological Sciences, 51(4):490–495, 1985.

Tadeusz J Kawecki and Dieter Ebert. Conceptual issues in local adaptation. Ecology letters, 7(12): 1225–1241, 2004.

Logan Kistler, S Yoshi Maezumi, Jonas Gregorio De Souza, Natalia AS Przelomska, Flaviane Malaquias Costa, Oliver Smith, Hope Loiselle, Jazmín Ramos-Madrigal, Nathan Wales, Eduardo Rivail Ribeiro, et al. Multiproxy evidence highlights a complex evolutionary legacy of maize in south america. Science, 362(6420):1309–1313, 2018.

Logan Kistler, Heather B Thakar, Amber M VanDerwarker, Alejandra Domic, Anders Bergström, Richard J George, Thomas K Harper, Robin G Allaby, Kenneth Hirth, and Douglas J Kennett. Archaeological central american maize genomes suggest ancient gene flow from south america. Proceedings of the National Academy of Sciences, 117(52):33124–33129, 2020.

Alexandra Kuznetsova, Per B Brockhoff, and Rune Haubo Bojesen Christensen. lmertest package: tests in linear mixed effects models. Journal of statistical software, 82(13), 2017.

Nick Lauter, Charles Gustus, Anna Westerbergh, and John Doebley. The inheritance and evolution of leaf pigmentation and pubescence in teosinte. Genetics, 167(4):1949–1959, 2004.

Roosa Leimu and Markus Fischer. A meta-analysis of local adaptation in plants. PloS one, 3(12): e4010, 2008.

Tuomas Leinonen, RJ Scott McCairns, Robert B O’hara, and Juha Merilä. Q st-f st comparisons: evolutionary and ecological insights from genomic heterogeneity. Nature Reviews Genetics, 14 (3):179–190, 2013.

Russell V Lenth. Using the lsmeans package. pain, 50(60):70, 2012.

Juul Limpens, Gustaf Granath, Rien Aerts, Monique MPD Heijmans, Lucy J Sheppard, Luca Bragazza, Berwyn L Williams, Håkan Rydin, Jill Bubier, Tim Moore, et al. Glasshouse vs field experiments: do they yield ecologically similar results for assessing n impacts on peat mosses? New Phytologist, 195(2):408–418, 2012.

Yann-Rong Lin, Keith F Schertz, and Andrew H Paterson. Comparative analysis of qtls affecting plant height and maturity across the poaceae, in reference to an interspecific sorghum population. Genetics, 141(1):391–411, 1995.

Dominique Louette and Melinda Smale. Farmers’ seed selection practices and traditional maize varieties in cuzalapa, mexico. Euphytica, 113(1):25–41, 2000.

Yoshihiro Matsuoka, Yves Vigouroux, Major M Goodman, Jesus Sanchez, Edward Buckler, and John Doebley. A single domestication for maize shown by multilocus microsatellite genotyping. Proceedings of the National Academy of Sciences, 99(9):6080–6084, 2002.

GG Medina, CJA Ruiz, and PRA Martínez. Los climas de méxico: una estratificación ambiental basada en el componente climático. Libro técnico, (1), 1998.

Kristin Mercer, Lesley Campbell, and Jing Luo. Effect of water availability and genetic diversity on flowering phenology, synchrony, and reproductive investment in maize. Maydica, 59(3):283–289, 2014.

Kristin L Mercer and Hugo Perales. Structure of local adaptation across the landscape: flowering time and fitness in mexican maize (zea mays l. subsp. mays) landraces. Genetic Resources and Crop Evolution, pages 1–19, 2018.

Kristin L Mercer and Hugo R Perales. Evolutionary response of landraces to climate change in centers of crop diversity. Evolutionary Applications, 3(5-6):480–493, 2010.

Kristin L Mercer, Ángel Martínez-Vásquez, and Hugo R Perales. Asymmetrical local adaptation of maize landraces along an altitudinal gradient. Evolutionary Applications, 1(3):489–500, 2008.

J Merilä and P Crnokrak. Comparison of genetic differentiation at marker loci and quantitative traits. Journal of Evolutionary Biology, 14(6):892–903, 2001.

William L Merrill, Robert J Hard, Jonathan B Mabry, Gayle J Fritz, Karen R Adams, John R Roney, and Art C MacWilliams. The diffusion of maize to the southwestern united states and its impact. Proceedings of the National Academy of Sciences, pages pnas–0906075106, 2009.

SM Mickelson, CS Stuber, L Senior, and SM Kaeppler. Quantitative trait loci controlling leaf and tassel traits in a b73× mo17 population of maize. Crop Science, 42(6):1902–1909, 2002.

Javier O Mijangos-Cortes, T Corona-Torres, D Espinosa-Victoria, A Muñoz-Orozco, J Romero-Peñaloza, and A Santacruz-Varela. Differentiation among maize (zea mays l.) landraces from the tarasca mountain chain, michoacan, mexico and the chalqueño complex. Genetic Resources and Crop Evolution, 54(2):309–325, 2007.

Ron Mittler. Abiotic stress, the field environment and stress combination. Trends in plant science, 11(1):15–19, 2006.

J Alberto Romero Navarro, Martha Willcox, Juan Burgueño, Cinta Romay, Kelly Swarts, Samuel Trachsel, Ernesto Preciado, Arturo Terron, Humberto Vallejo Delgado, Victor Vidal, et al. A study of allelic diversity underlying flowering-time adaptation in maize landraces. Nature genetics, 49(3):476, 2017.

Quetzalcóatl Orozco-Ramírez, Stephen B Brush, Mark N Grote, and Hugo Perales. A minor role for environmental adaptation in local-scale maize landrace distribution: results from a common garden experiment in oaxaca, mexico. Economic botany, 68(4):383–396, 2014.

R Ortega. Origen de la agricultura e importancia de los valles centrales de oaxaca. La Tecnología Agrícola Tradicional. Instituto Indigenísta Interamericano/Consejo Nacional de Ciencia y Tecnología, Oaxaca, México, pages 189–200, 1995.

Jason A Peiffer, Maria C Romay, Michael A Gore, Sherry A Flint-Garcia, Zhiwu Zhang, Mark J Millard, Candice AC Gardner, Michael D McMullen, James B Holland, Peter J Bradbury, et al. The genetic architecture of maize height. Genetics, 196(4):1337–1356, 2014.

H Perales. 23 landrace conservation of maize in mexico: an evolutionary breeding interpretation. Enhancing Crop Genepool Use: Capturing Wild Relative and Landrace Diversity for Crop Im-provement, page 271, 2016.

R Hugo Perales, Stephen B Brush, and Calvin O Qualset. Landraces of maize in central mexico: an altitudinal transect. Economic Botany, 57(1):7–20, 2003.

Kevin V Pixley, Gilberto E Salinas-Garcia, Anthony Hall, Martin Kropff, Cynthia Ortiz, Ankita Suhalia Bouvet, Prashant Vikram, and Sukhwinder Singh. Cimmyts seeds of discovery initiative: Harnessing biodiversity for food security and sustainable development. Indian J. Plant Genet. Resour, 30(3):231–240, 2017.

BM Prasanna et al. Phenotypic and molecular diversity of maize landraces: characterization and utilization. Indian J. Genet, 70(4):315–327, 2010.

Tanja Pyhäjärvi, Matthew B. Hufford, Sofiane Mezmouk, and Jeffrey Ross-Ibarra. Complex patterns of local adaptation in teosinte. Genome Biology and Evolution, 5(9):1594–1609, 07 2013.

R Core Team. R: A Language and Environment for Statistical Computing. R Foundation for Statistical Computing, Vienna, Austria, 2019. URL https://www.R-project.org/.

Peter Ranum, Juan Pablo Peña-Rosas, and Maria Nieves Garcia-Casal. Global maize production, utilization, and consumption. Annals of the New York Academy of Sciences, 1312(1):105–112, 2014.

Fausto Rodríguez-Zapata, Allison C Barnes, Karla A Blöcher-Juárez, Dan Gates, Andi Kur, Li Wang, Garrett M Janzen, Sarah Jensen, Juan M Estévez-Palmas, Taylor Crow, Rocío Aguilar-Rangel, Edgar Demesa-Arevalo, Tara Skopelitis, Sergio Pérez-Limón, Whitney L Stutts, Peter Thompson, Yu-Chun Chiu, David Jackson, Oliver Fiehn, Daniel Runcie, Edward S Buckler, Jeffrey Ross-Ibarra, Matthew B. Hufford, Ruairidh JH Sawers, and Rubén Rellán-Álvarez. Teosinte introgression modulates phosphatidylcholine levels and induces early maize flowering time. bioRxiv, 2021. doi: 10.1101/2021.01.25.426574. URL https://www.biorxiv.org/content/early/2021/01/26/2021.01.25.426574.

José Ariel Ruìz Corral, Noé Durán Puga, Jose de Jesus Sanchez Gonzalez, José Ron Parra, Diego Raymundo Gonzalez Eguiarte, JB Holland, and Guillermo Medina Garcia. Climatic adaptation and ecological descriptors of 42 mexican maize races. Crop Science, 48(4):1502–1512, 2008.

Guillermo Sarmiento. The dry plant formations of south america and their floristic connections. Journal of Biogeography, pages 233–251, 1975.

Outi Savolainen, Martin Lascoux, and Juha Merilä. Ecological genomics of local adaptation. Nature Reviews Genetics, 14(11):807, 2013.

PH Schuepp. Tansley review no. 59. leaf boundary layers. New Phytologist, pages 477–507, 1993.

Henry P Schwarcz and Margaret J Schoeninger. Stable isotope analyses in human nutritional ecology. American Journal of Physical Anthropology, 34(S13):283–321, 1991.

David A Selinger and Vicki L Chandler. Major recent and independent changes in levels and patterns of expression have occurred at the b gene, a regulatory locus in maize. Proceedings of the National Academy of Sciences, 96(26):15007–15012, 1999.

David A Selinger and Vicki L Chandler. B-bolivia, an allele of the maize b1 gene with variable expression, contains a high copy retrotransposon-related sequence immediately upstream. Plant Physiology, 125(3):1363–1379, 2001.

Bekele Shiferaw, Boddupalli M Prasanna, Jonathan Hellin, and Marianne Bänziger. Crops that feed the world 6. past successes and future challenges to the role played by maize in global food security. Food Security, 3(3):307, 2011.

David B Stern, Maureen R Hanson, and Alice Barkan. Genetics and genomics of chloroplast biogenesis: maize as a model system. Trends in plant science, 9(6):293–301, 2004.

Shohei Takuno, Peter Ralph, Kelly Swarts, Rob J Elshire, Jeffrey C Glaubitz, Edward S Buckler, Matthew B Hufford, and Jeffrey Ross-Ibarra. Independent molecular basis of convergent highland adaptation in maize. Genetics, pages genetics-115, 2015.

Maud Irène Tenaillon and Alain Charcosset. A european perspective on maize history. Comptes rendus biologies, 334(3):221–228, 2011.

Göte Turesson. The genotypical response of the plant species to the habitat. Hereditas, 3(3): 211–350, 1922.

Robert J Twohey III, Lucas M Roberts, and Anthony J Studer. Leaf stable carbon isotope composition reflects transpiration efficiency in zea mays. The Plant Journal, 97(3):475–484, 2019.

Joost Van Heerwaarden, John Doebley, William H Briggs, Jeffrey C Glaubitz, Major M Goodman, Jose de Jesus Sanchez Gonzalez, and Jeffrey Ross-Ibarra. Genetic signals of origin, spread, and introgression in a large sample of maize landraces. Proceedings of the National Academy of Sciences, 108(3):1088–1092, 2011.

Sandy Vanderauwera, Philip Zimmermann, Stéphane Rombauts, Steven Vandenabeele, Christian Langebartels, Wilhelm Gruissem, Dirk Inzie, and Frank Van Breusegem. Genome-wide analysis of hydrogen peroxide-regulated gene expression in arabidopsis reveals a high light-induced transcriptional cluster involved in anthocyanin biosynthesis. Plant Physiology, 139(2):806–821, 2005.

Tania Carolina Camacho Villa, Nigel Maxted, Maria Scholten, and Brian Ford-Lloyd. Defining and identifying crop landraces. Plant Genetic Resources, 3(3):373–384, 2005.

Li Wang, Timothy M Beissinger, Anne Lorant, Claudia Ross-Ibarra, Jeffrey Ross-Ibarra, and Matthew B Hufford. The interplay of demography and selection during maize domestication and expansion. Genome biology, 18(1):215, 2017.

Li Wang, Emily B Josephs, Kristin M Lee, Lucas M Roberts, Rubén Rellán-Álvarez, Jeffrey Ross-Ibarra, and Matthew B Hufford. Molecular parallelism underlies convergent highland adaptation of maize landraces. bioRxiv, 2020.

Jennifer Watling, Myrtle P Shock, Guilherme Z Mongeló, Fernando O Almeida, Thiago Kater, Paulo E De Oliveira, and Eduardo G Neves. Direct archaeological evidence for southwestern amazonia as an early plant domestication and food production centre. PLoS One, 13(7): e0199868, 2018.

Hernandez E Wellhausen EJ, Roberts LM. Races of Maize in Mexico. The Bussey Institution, Harvard University Press, Cambridge, MA, 1952.

Peter Wenzl, Jason Carling, David Kudrna, Damian Jaccoud, Eric Huttner, Andris Kleinhofs, and Andrzej Kilian. Diversity arrays technology (dart) for whole-genome profiling of barley. Proceedings of the National Academy of Sciences, 101(26):9915–9920, 2004.

Michael C Whitlock. Evolutionary inference from qst. Molecular ecology, 17(8):1885–1896, 2008.

Hong Xu, Tracy E Twine, and Evan Girvetz. Climate change and maize yield in iowa. PLoS One, 11(5):e0156083, 2016.

